# Acetylcholine prioritises direct synaptic inputs from entorhinal cortex to CA1 by differential modulation of feedforward inhibitory circuits

**DOI:** 10.1101/2020.01.20.912873

**Authors:** Jon Palacios-Filardo, Matt Udakis, Giles A. Brown, Benjamin G. Tehan, Miles S. Congreve, Pradeep J. Nathan, Alastair J.H. Brown, Jack R. Mellor

## Abstract

Acetylcholine release in the hippocampus plays a central role in the formation of new memory representations by facilitating synaptic plasticity. It is also proposed that memory formation requires acetylcholine to enhance responses in CA1 to new sensory information from entorhinal cortex whilst depressing inputs from previously encoded representations in CA3, but this influential theory has not been directly tested. Here, we show that excitatory inputs from entorhinal cortex and CA3 are depressed equally by synaptic release of acetylcholine in CA1. However, greater depression of feedforward inhibition from entorhinal cortex results in an overall enhancement of excitatory-inhibitory balance and CA1 activation. Underpinning the prioritisation of entorhinal inputs, entorhinal and CA3 pathways engage distinct feedforward interneuron subpopulations and depression is mediated differentially by presynaptic muscarinic M_3_ and M_4_ receptors respectively. These mechanisms enable acetylcholine to prioritise novel information inputs to CA1 during memory formation and suggest selective muscarinic targets for therapeutic intervention.

## Introduction

Cognitive processing in the brain must continuously adapt to changing environmental situations. However, the stability of physical connectivity within neuronal networks, at least over relatively short timescales (< min), means that the brain requires systems that can enact rapid functional network reconfigurations. Release of neuromodulator transmitters via long-range projections fulfils the requirements for functional reconfiguration (Marder, 2012) and occurs in response to situations that demand behavioural or cognitive adaptation (Dayan, 2012). But the mechanisms by which neuromodulators such as acetylcholine reconfigure neuronal networks remain largely unknown.

The widespread release of acetylcholine within the brain is historically associated with arousal and attention (Hasselmo and Sarter, 2011; Micheau and Marighetto, 2011; Robbins, 1997; Teles-Grilo Ruivo and Mellor, 2013). More recently it has also been found to be associated with unexpected rewards or punishments (Hangya and Kepecs, 2015; Teles-Grilo Ruivo et al., 2017) signalling the need to update existing representations with new salient information. To achieve this acetylcholine must reconfigure neural networks in two key ways: (i) open a window for encoding new memories or updating existing ones, and (ii) prioritise new sensory information for incorporation into memory ensembles (Hasselmo, 2006; Palacios-Filardo and Mellor, 2019). Acetylcholine facilitates the induction of synaptic plasticity thereby opening a window for the creation of memory ensembles (Buchanan et al., 2010; Dennis et al., 2016; Hasselmo et al., 1995; Isaac et al., 2009; Mitsushima et al., 2013; Shinoe et al., 2005) and it increases the output gain from primary sensory cortices enhancing signal-to-noise for new sensory information (Eggermann et al., 2014; Fu et al., 2014; Letzkus et al., 2011). It is also proposed to prioritise sensory inputs from the neocortex into memory ensembles within the hippocampus (Dannenberg et al., 2017; Hasselmo, 2006; Hasselmo and Schnell, 1994; Hasselmo et al., 1995) but this critical component of the mechanism by which acetylcholine gates the updating of memory representations has yet to be tested in detail.

The hippocampus is a hub for the encoding, updating and retrieval of episodic memories, enabling events to be placed into a context. Individual items of information from the neocortex are thought to be sparsely encoded and separated by strong lateral inhibition in the dentate gyrus before being assembled into larger memory representations within the recurrent CA3 network (Hasselmo, 2006; Prince et al., 2016). These memory representations are then transferred via the Schaffer collateral (SC) pathway to CA1 which also receives new sensory information directly from the entorhinal cortex layer III pyramidal neurons via the temporoammonic (TA) pathway enabling CA1 to compare and integrate the new information (Ahmed and Mehta, 2009; Eichenbaum, 2017; Takahashi and Magee, 2009; Witter, 1993). It is therefore predicted that acetylcholine enhances the relative weights of TA inputs to CA1 over SC inputs during memory formation.

Perhaps counter-intuitively, acetylcholine inhibits both TA and SC glutamatergic excitatory transmission in CA1. In the SC pathway this occurs via presynaptic muscarinic M_4_ receptors but the identity of the receptors mediating depression at the TA pathway is unclear (Dasari and Gulledge, 2011; Goswamee and McQuiston, 2019; Thorn et al., 2017). The anatomically segregated targeting of TA and SC inputs to distal and more proximal dendritic locations on CA1 pyramidal neurons respectively (Witter, 1993) together with muscarinic receptor specificity provide potential mechanisms for differential sensitivity to acetylcholine and therefore altering the relative weights of synaptic input. However, the evidence for this is equivocal with exogenously applied cholinergic agonists indicating that SC transmission is more sensitive to cholinergic modulation than TA transmission (Hasselmo and Schnell, 1994) but the reverse reported for endogenous synaptically released acetylcholine (Goswamee and McQuiston, 2019).

An alternative mechanism by which acetylcholine might rebalance the relative weight s of SC and TA inputs is the modulation of the intrinsic and synaptic properties of hippocampal GABAergic interneurons (Cea-del Rio et al., 2011; Cea-del Rio et al., 2010; Leao et al., 2012; Szabo et al., 2010) which have a profound impact on CA1 pyramidal neuron input integration rules and subsequent output (Leao et al., 2012; Milstein et al., 2015). Feedforward interneurons in the SC pathway are primarily perisomatic targeting basket cells expressing parvalbumin (PV) or cholecystokinin (CCK) (Basu et al., 2013; Freund and Katona, 2007; Glickfeld and Scanziani, 2006; Klausberger and Somogyi, 2008; Milstein et al., 2015) whose inhibition is strongly regulated by acetylcholine (Cea-del Rio et al., 2011; Szabo et al., 2010) whereas the mediators of feedforward inhibition in the TA pathway are primarily CCK or neuropeptide Y (NPY) expressing interneurons (Basu et al., 2013; Klausberger and Somogyi, 2008; Milstein et al., 2015) that are also potentially regulated by acetylcholine (Cea-del Rio et al., 2010; Raza et al., 2017). Moreover, feedback inhibition via oriens lacunosum moleculare (OLM) interneurons, which specifically target the same distal dendritic regions as the TA pathway, are directly excited by acetylcholine (Leao et al., 2012; Pouille and Scanziani, 2004). This indicates that cholinergic modulation of inhibition within the hippocampal circuit strongly dictates excitatory input integration and CA1 output, but the integrated effect of acetylcholine on the hippocampal network and its input-output function has not been investigated.

In this study we tested the hypothesis that acetylcholine release in the hippocampus prioritises new sensory input to CA1 via the TA pathway over internal representations via the SC pathway. We find that endogenous synaptically released acetylcholine depresses SC and TA excitatory inputs equally but that feedforward inhibition in the TA pathway is more sensitive to cholinergic modulation. This produces an increase in excitatory-inhibitory ratio selectively for the TA pathway driven by differential regulation of interneuron subpopulations and distinct muscarinic receptor subtypes. We therefore provide a mechanism by which acetylcholine dynamically prioritises sensory information direct from entorhinal cortex over internal representations held in CA3.

## Results

### Endogenous acetylcholine release modulates synaptic inputs to CA1

To enable selective activation of endogenous acetylcholine release we expressed the light-activated cation channel channelrhodopsin-2 (ChR2) in a cre-dependent manner using mice that express cre recombinase under control of the promoter for Choline AcetylTransferase (ChAT-cre) crossed with mice expressing cre-dependent ChR2 (ChAT-ChR2 mice; methods). Immunohistochemisty confirmed that ChR2 was expressed in cholinergic cells within the medial septum (Figure 1A-B) whose axon fibers densely innervated the dorsal hippocampus (Figure 1C) in agreement with the previously described anatomy (Teles-Grilo Ruivo and Mellor, 2013). Whole-cell patch clamp recordings from medial septal neurons expressing ChR2 confirmed they fired action potentials in response to 5ms of 470nm light up to a maximum frequency of ∼25Hz (Figure 1B). We also confirmed that light stimulation in hippocampal slices resulted in acetylcholine release. Recordings from interneurons located in stratum oriens revealed fast synaptic responses to light stimulation mediated by nicotinic receptors (Figure 1D) consistent with activation of cholinergic axons and endogenous release of acetylcholine (Leao et al., 2012). In these recordings and further recordings from CA1 pyramidal cells we saw no inhibitory postsynaptic currents that might be caused by light-evoked co-release of GABA or glutamate from either local or long-range ChAT expressing neurons (Figure 1E) (Takacs et al., 2018; Yi et al., 2015).

**Figure 1.**
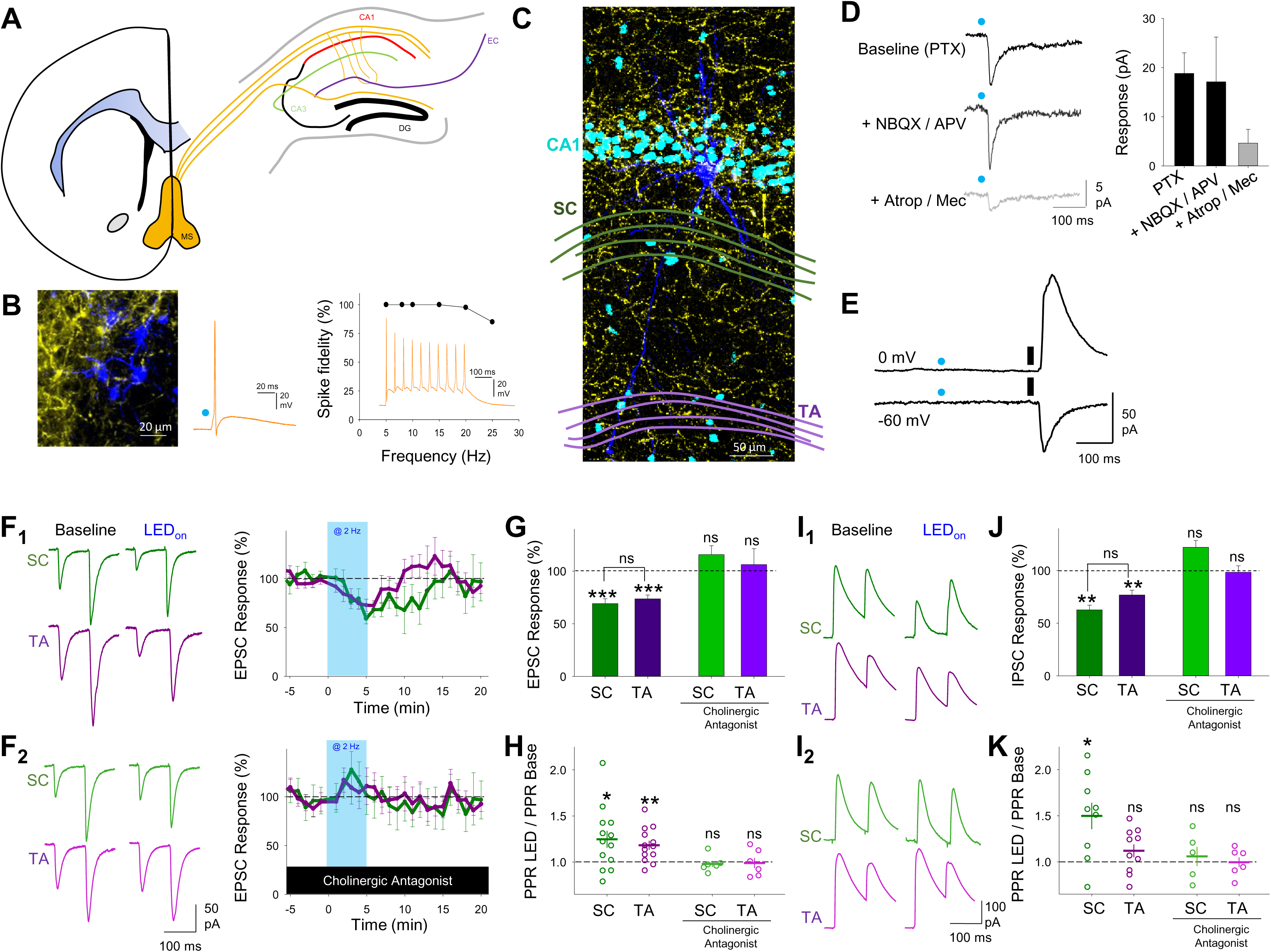
Endogenous release of acetylcholine reduces excitatory and inhibitory synaptic inputs to CA1 pyramidal neurons. **A,** Coronal section illustration of medial septum (MS, yellow) and its projections to dorsal hippocampus including Schaffer collateral (green) and temporoammonic (purple) inputs to CA1 from CA3 and entorhinal cortex (EC) respectively. **B**, Immunofluorescence of cholinergic neurons in medial septum filled with neurobiotin (blue) expressing ChR2-YFP protein (left) and light evoked stimulation (blue dot, 10ms) of cholinergic neuron, which reliably elicits action potentials at frequencies < 25 Hz (right). **C**, Immunofluorescence of CA1 area of the hippocampus highlighting a CA1 pyramidal neuron filled with neurobiotin (blue) and surrounding cholinergic axons (yellow). Nuclei stained with DAPI (light blue) and location of Schaffer collateral (SC) and temporoammonic (TA) axons illustrated in green and purple respectively. **D**, Light-evoked stimulation of cholinergic axons (blue dot) elicits fast synaptic responses in Stratum Oriens interneurons recorded at −60 mV that are sensitive to atropine (25 µM) and mecamylamine (50 µM) but not picrotoxin (PTX, 50 µM), NBQX (20 µM) or D-APV (25 µM). **E,** No response to light-evoked stimulation of cholinergic axons (blue dot) was seen in CA1 pyramidal neurons recorded at 0 mV or −60 mV in contrast to electrical stimulation (black line). **F**, SC (green) and TA (purple) evoked EPSCs in CA1 pyramidal neurons are reversibly depressed by endogenous release of acetylcholine evoked by 5 minutes light stimulation at 2 Hz (F_1_). The depression of EPSCs is blocked by application of cholinergic antagonists atropine (25 µM) and mecamylamine (50 µM) (F_2_). **G-H**, Acetylcholine release depressed SC and TA pathway evoked EPSCs (G) and increased paired-pulse ratio (H, PPR). **I**, Feedforward disynaptic IPSCs evoked by stimulation of SC and TA pathways are depressed by light evoked acetylcholine release (I_1_). The depression of IPSCs is blocked by application of cholinergic antagonists atropine and mecamylamine (I_2_). **J-K**, Effects of acetylcholine release on SC and TA pathway evoked IPSC response (J) and PPR (K). Data are mean ± SEM; Two-tailed paired Student’s T-test *** p < 0.001 ** p < 0.01 * p < 0.05; g-h and j-k inter group comparison one-way ANOVA with post hoc Bonferroni correction.

To selectively activate the Schaffer collateral and temporoammonic pathways into CA1 stimulating electrodes were placed within the two axon pathways in dorsal hippocampal slices. This enabled independent stimulation of each pathway and the engagement of both the direct excitatory inputs and disynaptic feedforward inhibitory inputs without activating direct inhibitory inputs, demonstrated by the blockade of inhibitory inputs by NBQX (20 µM) (Figure S1A-B). We also pharmacologically confirmed the identity of the TA input by application of the mGluR group II/III agonist DCG-IV (3 µM) that selectively inhibits glutamate release from temporoammonic pathway terminals (Ceolin et al., 2011) (Figure S1C).

To test the effect of endogenous acetylcholine release on synaptic inputs to CA1, hippocampal slices were stimulated with light at a frequency of 2 Hz for 5 minutes to evoke physiologically maximal acetylcholine release (Jing et al., 2018). In the presence of the GABA_A_ receptor antagonist picrotoxin, isolated SC and TA pathway excitatory postsynaptic current (EPSC) amplitudes were depressed by very similar amounts (Figure 1F-G; SC pathway 69 ± 5 %, n = 12 from 6 mice, p < 0.001_;_ TA pathway 74 ± 4 %, n = 13 from 6 mice, p < 0.001) with a concomitant increase in the paired-pulse ratio (PPR) (Figure 1H; SC pathway, 125 ± 9 %, p < 0.05; TA pathway, 118 ± 5 %, p < 0.01), indicating a presynaptic locus of action. Application of nicotinic and muscarinic receptor antagonists atropine (25 µM) and mecamylamine (50 µM) blocked the effects of endogenous acetylcholine release (Figure 1F-H; SC pathway 116 ± 8 %, n = 6 from 3 mice, p > 0.05_;_ TA pathway 106 ± 15 %, n = 6 from 3 mice, p > 0.05). Therefore, contrary to our initial hypothesis (Dannenberg et al., 2017; Goswamee and McQuiston, 2019; Hasselmo, 2006; Hasselmo and Schnell, 1994), acetylcholine did not inhibit one pathway more than the other but instead depressed both equally.

We next tested disynaptic feedforward inhibitory postsynaptic currents (IPSCs) in response to stimulation of SC or TA pathways. The amplitude of evoked IPSCs was also reduced by endogenous acetylcholine release (Figure 1I-J; SC pathway, 63 ± 5 %, n = 9 from 6 mice, p < 0.01; TA pathway, 77 ± 5 %, n = 10 from 6 mice, p < 0.01) but surprisingly IPSC PPR was only increased in the SC pathway (Figure 1I,K; SC pathway, 150 ± 14 %, p < 0.05; TA pathway, 112 ± 7 %, p > 0.05). Similar to EPSCs, the reduction in IPSCs was completely blocked by muscarinic and nicotinic receptor antagonists (Figure 1I-K; SC pathway IPSC 122 ± 6 % and PPR 106 ± 10 %, n = 5 from 4 mice, p > 0.05; TA pathway IPSC 98 ± 6 % and PPR 99 ± 6 %, n = 6 from 4 mice, p > 0.05). The observation that IPSCs were depressed equally in each pathway but PPR was increased in the SC pathway suggests that during repetitive stimulation inhibitory drive will increase in the SC pathway relative to the TA pathway. This predicts that although acetylcholine depresses excitatory synaptic transmission in the TA and SC pathways equally, its overall effect on excitatory-inhibitory ratio favours TA inputs during repetitive stimulation when the effects of acetylcholine on feedforward inhibition are taken into account.

### Differential cholinergic modulation of excitatory-inhibitory ratio for Schaffer collateral and temporoammonic inputs to CA1

To test whether excitatory-inhibitory balance was differentially altered between SC and TA input pathways we recorded monosynaptic EPSCs and disynaptic feedforward IPSCs for SC and TA pathways in the same CA1 pyramidal neuron (see methods; Figure 2A). 5 consecutive stimuli at 10 Hz were given alternately to SC then TA pathway to determine the evolution of synaptic modulation by acetylcholine during a repetitive train of stimuli. In these experiments we mimicked the release of endogenous acetylcholine with application of the cholinergic receptor agonist carbachol (CCh), a non-hydrolysable analogue of acetylcholine that is not selective between cholinergic receptor subtypes. Application of increasing concentrations of CCh revealed that 10 µM CCh was required to induce depression for both EPSCs and IPSCs in both SC and TA pathways similar to endogenous acetylcholine release (Figure S2A; SC pathway EPSC, 35 ± 6 %, n = 20 from 11 mice; TA pathway EPSC, 50 ± 5 %, n = 20 from 11 mice; SC pathway IPSC, 29 ± 3 %, n = 20 from 11 mice; TA pathway IPSC, 40 ± 4 %, n = 20 from 11 mice), but at lower concentrations of CCh SC excitatory synaptic transmission showed higher sensitivity to CCh than the TA pathway (Hasselmo and Schnell, 1994) suggestive of different receptor affinities or signalling pathways regulating presynaptic release (Figure S2A; CCh 1 µM at SC pathway, 52 ± 6 %, n = 9 from 4 mice; TA pathway, 91 ± 18 %, n = 9 from 4 mice). The depression of EPSCs and IPSCs with 10 µM CCh occurred for all responses in both SC and TA pathways (Figure 2A and Figure S2B) but the degree of depression was not consistent between pathways over the course of repetitive stimulation. Cholinergic receptor activation enhanced synaptic facilitation and increased PPR for excitatory and feedforward inhibitory connections in the SC pathway, while the TA pathway only displayed a marked increase in PPR in excitatory but not feedforward inhibitory inputs (Figure 2B and Figure S2B-E; 5^th^ stimuli PPR change for SC EPSC, 197 ± 23 %, p < 0.01; SC IPSC, 188 ± 13 %, p < 0.001; TA EPSC, 170 ± 13 %, p < 0.001; TA IPSC, 120 ± 13 %, p > 0.05, n = 20 from 11 mice), supporting the initial results using endogenous acetylcholine release. Indeed, the close similarity in PPR increase for both excitatory and feedforward inhibitory transmission in the SC pathway ensured that the excitatory-inhibitory (E-I) ratio in the SC pathway did not change after cholinergic receptor activation for any stimuli within the train (Figure 2C; 5^th^ stimuli on SC E-I ratio, 0.29 ± 0.05 and 0.41 ± 0.10, for baseline and CCh respectively, p > 0.05). Conversely, excitation-inhibition ratio in the TA pathway showed a marked increase after CCh application that evolved over the course of the train of stimuli (Figure 2C; 5^th^ stimuli on TA E-I ratio, 0.34 ± 0.06 and 0.6 ± 0.10, for baseline and CCh respectively, p < 0.001). This meant that over the course of the train the TA input exerted relatively greater influence over the postsynaptic neuron compared to the SC input when cholinergic receptors were activated, as demonstrated by the comparison of excitation-inhibition ratio between the SC and TA pathways (Figure 2D; 5^th^ stimuli on TA/SC E-I ratio, 1.28 ± 0.25 and 1.7 ± 0.25, for baseline and CCh respectively, p < 0.01). These data show that differential modulation of feedforward inhibition between SC and TA pathways by cholinergic receptor activation produces an increase in the relative strength of the TA input to CA1 pyramidal neurons. Furthermore, the data suggest that SC and TA pathways engage distinct local inhibitory interneuron populations with different overall short-term dynamic responses to acetylcholine.

**Figure 2.**
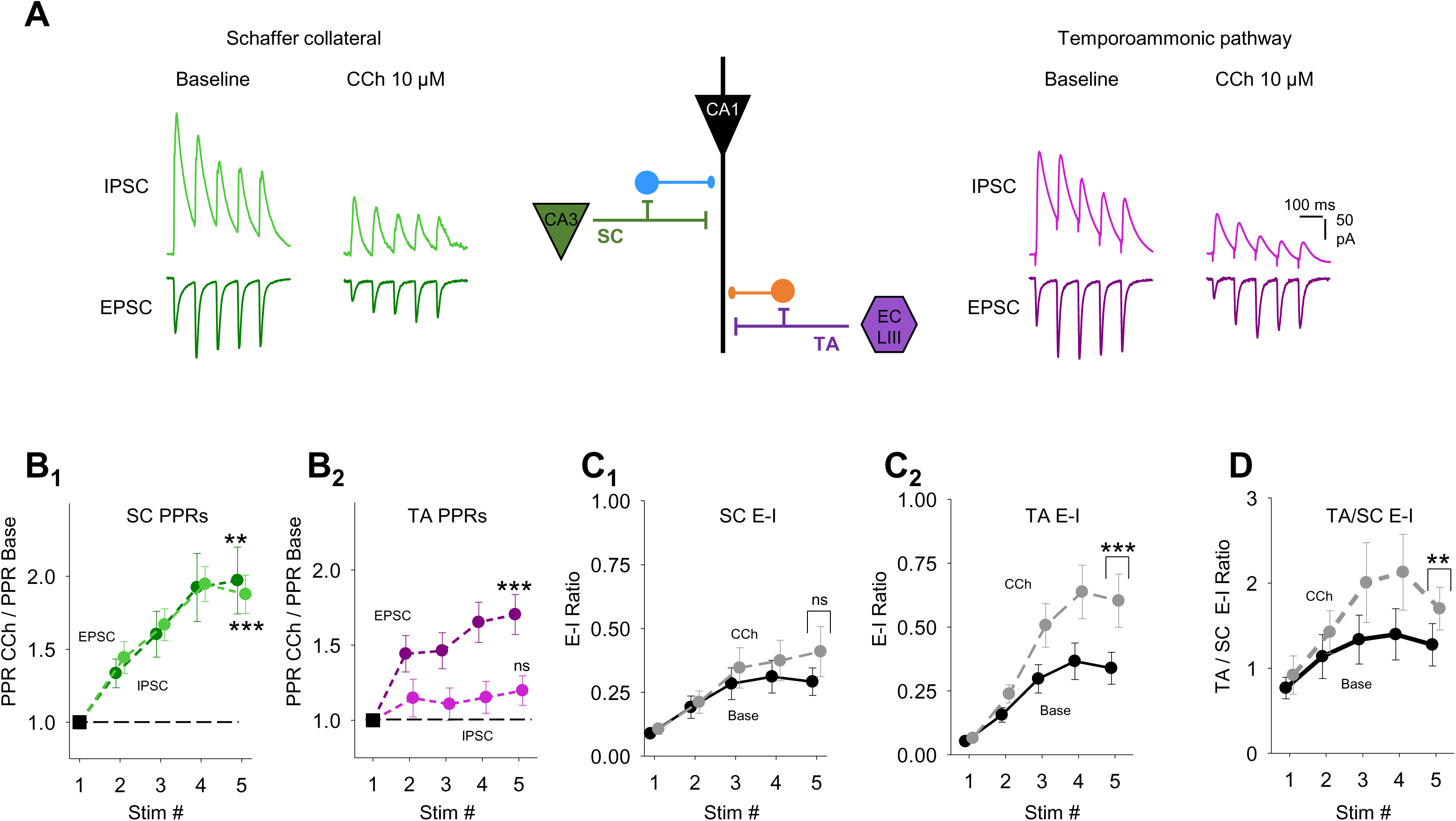
Cholinergic receptor activation enhances excitatory-inhibitory balance for temporoammonic synaptic inputs relative to Schaffer collateral inputs. **A**, Middle, schematic representation of the experimental approach incorporating simultaneous recording of excitatory (V_h_ = −60 mV) and feedforward inhibitory (V_h_ = 0 mV) synaptic inputs from Schaffer collateral (SC) and temporoammonic (TA) input pathways to CA1 pyramidal neuron (bottom). Example traces for EPSCs and IPSCs in response to trains of 5 stimuli at 10 Hz to SC (green, left) and TA (purple, right) pathways before and after carbachol (CCh, 10µM) application. **B**, Change in paired-pulse ratio (PPR) after CCh application for excitatory and inhibitory responses to SC (B_1_) and TA (B_2_) pathway stimulation. PPR is measured compared to the first response for each response in the train. **C**, Comparison of synaptic Excitatory-Inhibitory (E-I) ratio before and after CCh application measured by charge transfer at Vh = −60 mV and 0 mV for SC (C_1_) and TA (C_2_) input pathways. **D**, Comparison of synaptic E-I ratio between TA and SC input pathways before and after CCh application. CCh enhanced the overall relative synaptic charge transfer from TA pathway. Data are mean ± SEM; Two tailed Student’s paired T-test *** p < 0.001 ** p < 0.01 * p < 0.05.

### Cholinergic modulation of interneuron recruitment to feedforward inhibitory synaptic transmission

Feedforward interneurons in the SC pathway are primarily perisomatic targeting basket cells expressing parvalbumin (PV^+^) or cholecystokinin (CCK^+^) whereas the mediators of feedforward inhibition in the TA pathway are likely dendritically targeting CCK^+^ or neuropeptide Y (NPY^+^) expressing interneurons (Basu et al., 2013; Freund and Katona, 2007; Glickfeld and Scanziani, 2006; Klausberger and Somogyi, 2008; Milstein et al., 2015). Analysis of our recordings revealed that feedforward SC IPSCs had faster decay kinetics than TA IPSCs (Figure 3A-C; SC IPSC decay tau, 43.0 ± 2.7 ms, n = 45 from 24 mice vs TA IPSC decay tau, 60.1 ± 3.4 ms, n = 92 from 36 mice, p < 0.005) in accordance with predictions that the more distal synaptic location of inhibitory inputs from TA feedforward interneurons and therefore increased dendritic filtering means that these IPSCs have slower kinetics (Milstein et al., 2015). GABAergic synapses from PV^+^ and NPY^+^, but not CCK^+^, interneurons onto CA1 pyramidal cells are depressed by µ-opioid receptors (Glickfeld et al., 2008; Gulyas et al., 2010; Krook-Magnuson et al., 2011). SC IPSCs were more sensitive to µ-opioid receptor agonist DAMGO (1 µM) than TA IPSCs (Figure 3D-E; IPSC 1^st^ response peak after DAMGO, 51 ± 4 % and 69 ± 5 %, for SC and TA respectively, n = 11 from 4 mice, p < 0.05) indicating that in our experiments PV^+^ interneurons form a major component of feedforward inhibition in the SC pathway whereas CCK^+^ interneurons form the major component of feedforward inhibition in the TA pathway. There are also minor components from other interneuron subtypes, most likely CCK^+^ basket cells in the SC pathway and PV^+^ or NPY^+^ interneurons in the TA pathway (Basu et al., 2013; Freund and Katona, 2007; Glickfeld and Scanziani, 2006; Klausberger and Somogyi, 2008; Milstein et al., 2015).

**Figure 3.**
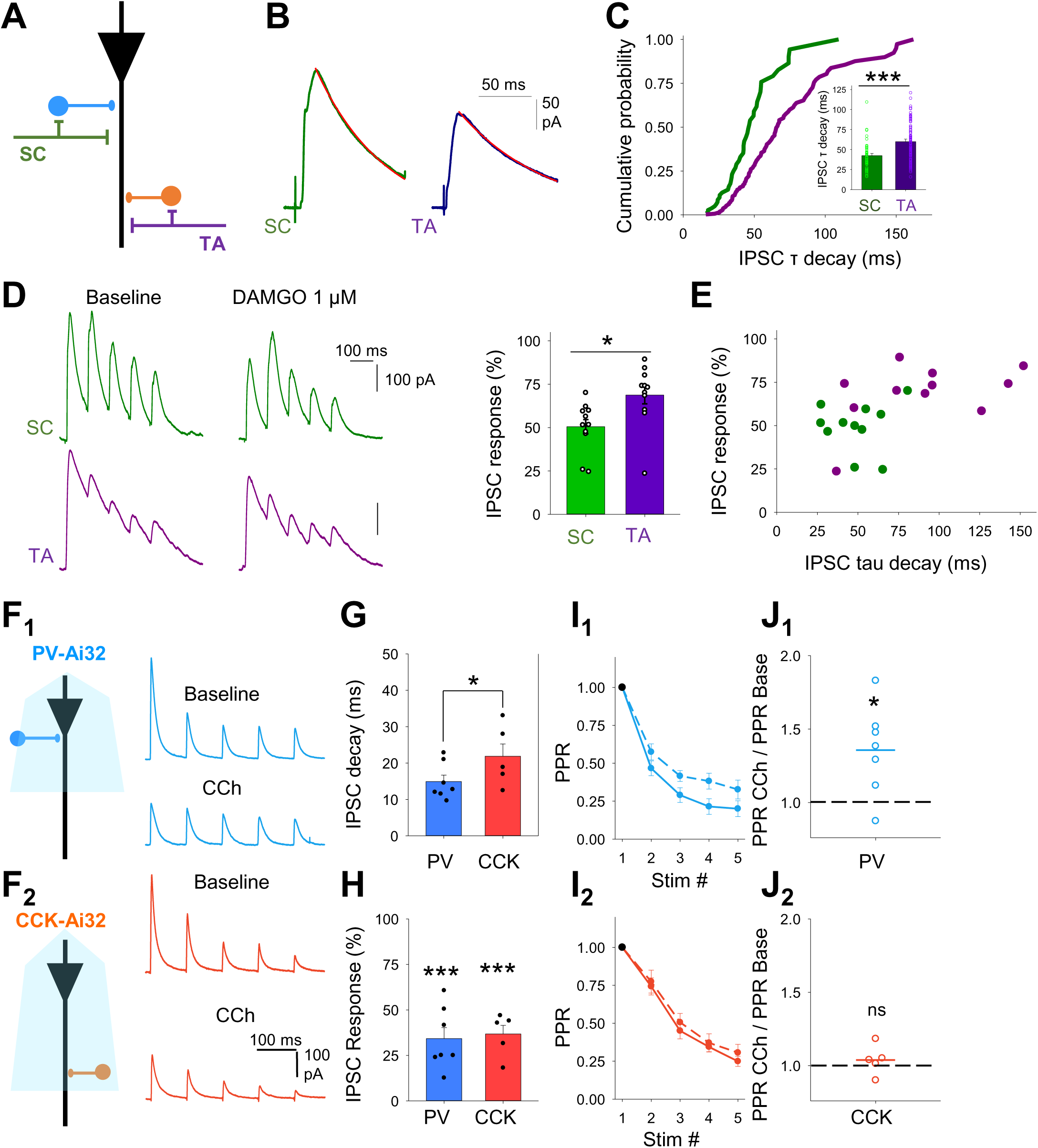
Cholinergic modulation of inhibitory inputs from distinct feedforward interneuron populations. **A**, Schematic representation of different feedforward interneuron populations engaged by Schaffer collateral (SC) and temporoammonic (TA) pathways within CA1. **B-C**, Disynaptic feedforward IPSCs (B) and distribution of decay kinetics (C) for Schaffer collateral (SC, green) and temporoammonic (TA, purple) input pathways demonstrating distinct populations of feedforward interneurons. Quantification of the IPSC tau decay (insert C). **D**, µ-opioid receptor agonist DAMGO (1 µM) depression of disynaptic feedforward IPSCs from SC and TA pathways. **E**, IPSC decay kinetics and sensitivity to DAMGO correlate and distinguish SC from TA evoked IPSCs. **F,** Optogenetic activation of either PV (F**_1_**) or CCK (F**_2_**) interneurons at 10 Hz evoked a train of IPSCs in CA1 pyramidal neurons. IPSCs from both interneurons are depressed by CCh (10 µM). **G-H**. IPSCs from PV interneurons display faster decay kinetics than IPSCs from CCK interneurons (G) but CCh depressed the IPSC amplitudes of the first responses in the train by a similar amount (H). **I-J**. IPSCs from both PV and CCK interneurons demonstrated frequency-dependent depression. Frequency-dependent depression was reduced after CCh application for PV (I_1_-J_1_) but not CCK (I_2_-J_2_) evoked IPSCs. Data are mean ± SEM; Two tailed Student’s paired T-test *** p < 0.001 ** p < 0.01 * p < 0.05.

The engagement of different interneuron subtypes in feedforward inhibition in the SC and TA pathways might explain the differential modulation of feedforward inhibition by acetylcholine. Therefore, we investigated whether the output from these interneurons onto CA1 pyramidal cells is modulated by acetylcholine and, if so, whether modulation evolves differentially for the 2 inputs during a burst of responses. To test this we used mice expressing ChR2 in PV^+^ or CCK^+^ interneurons (see methods) and gave a train of 5 light stimuli at 10 Hz to the slices whilst recording IPSCs from pyramidal neurons at 0 mV in the presence of NBQX and D-APV to avoid recording glutamatergic, disynaptic inhibitory inputs or ChR2 currents (Figure S3). To test the sensitivity of PV^+^ and CCK^+^ synapses to cholinergic modulation, CCh was bath applied to the slice whilst selectively evoking either PV^+^ or CCK^+^ derived IPSCs (Figure 3F). PV^+^ evoked IPSCs displayed faster decay kinetics to CCK^+^ evoked IPSCs supporting their perisomatic and dendritic synaptic locations respectively (Figure 3G; PV^+^ decay kinetics, 14.9 ± 1.8 ms, n = 7 vs CCK^+^ decay kinetics, 21.9 ± 3.4 ms, n = 5, p < 0.05). Decay kinetics of optogenetically evoked IPSCs were faster than disynaptically evoked feedforward IPSCs as predicted for inputs with greater synchrony. CCh depressed IPSCs from both PV^+^ and CCK^+^ synapses indicating a direct cholinergic modulation of these interneurons (Figure 3F,H; PV^+^ responses, 34.3 ± 6.0 %, n = 7, p < 0.005; CCK^+^ responses, 37.8 ± 4.8 %, n = 5, p < 0.005). Both synapses exhibited frequency-dependent depression but CCh selectively increased PPR of PV^+^ but not CCK^+^ synapses (Figure 3I-J; PV^+^ IPSC PPR, 136 ± 11 %, n = 7, p < 0.05; CCK^+^ IPSC PPR, 104 ± 4 %, n = 5, p > 0.05). The lack of effect of cholinergic receptor activation on PPR at CCK^+^ synapses mirrors the lack of effect on PPR for feedforward inhibition in the TA pathway and confirms that CCK^+^ interneurons are the major component of feedforward inhibition in the TA pathway whereas PV^+^ interneurons and synapses that increase PPR form feedforward inhibition in the SC pathway. The differential effect of acetylcholine at PV^+^ and CCK^+^ synapses provides a mechanism for the enhancement of TA pathway excitatory-inhibitory ratio in comparison to SC pathway.

### Presynaptic muscarinic M_3_ receptor modulation of TA pathway excitatory and feedforward inhibitory synaptic transmission

The synaptic depression of Schaffer collateral inputs to CA1 by acetylcholine is characterised genetically and pharmacologically to be mediated by muscarinic M_4_ receptors (Dasari and Gulledge, 2011; Thorn et al., 2017). This was confirmed by application of the dual muscarinic M_4_ and M_1_ receptor agonist compond1 (1 µM; Figure S4), which selectively depressed SC but not TA pathway excitatory inputs (Figure 4A-C; SC EPSC response, 63 ± 5 %, n = 17, from 8 mice, p < 0.001; TA EPSC response, 94 ± 5 %, n = 17, from 8 mice, p > 0.05). However, the identity of cholinergic receptors mediating the depression of TA inputs is unclear. Therefore, we aimed to determine which cholinergic receptors modulate TA pathway feedforward excitatory and inhibitory synaptic transmission onto CA1 pyramidal neurons. TA pathway excitatory synaptic transmission was isolated by recording in the presence of PTX and holding the membrane voltage at −65 mV (see methods; Figure 4D). Similar to previous results (Figures 1&2), TA EPSCs were depressed by application of 10 µM CCh and PPR was increased (Figure 4E; EPSC response, 45 ± 3 %, n = 10 from 5 mice, p < 0.01; PPR, 129 ± 8 %, n = 10 from 5 mice, p < 0.05). These data suggest a presynaptic locus of action of cholinergic receptors. We next pharmacologically dissected which cholinergic receptor subtypes were involved. Application of the non-selective nicotinic receptor antagonist mecamylamine (25 µM) had no effect on CCh depression of EPSCs (Figure 4F; 40.6 ± 9.5 %, n = 6 from 3 mice, p < 0.01) and PPR (Figure 4G; 124 ± 8 %, n = 6 from 3 mice, p < 0.05), while the non-selective muscarinic receptor antagonist atropine (10 µM) blocked the decrease of EPSCs (Figure 4F; 91 ± 4 %, n = 6 from 3 mice, p > 0.05) and prevented the increase in PPR (Figure 4G; 105 ± 4 %, n = 6 from 23 mice, p > 0.05), suggesting a direct involvement of muscarinic receptors. Muscarinic M_1_ receptor agonist GSK-5 (500 nM) (Dennis et al., 2016) did not replicate CCh depression of EPSCs and increase in PPR (Figure 4F-G; EPSCs, 91 % ± 4 %, PPR 101 ± 5 %, n = 7 from 4 mice, p > 0.05) nor did the selective M_1_ receptor antagonist, nitrocaramiphen (100 nM) prevent CCh induced depression and increase in PPR (Figure 4F-G; EPSC 51 ± 4 %, n = 6 from 4 mice, p < 0.01; PPR 124 ± 6 %, n = 6 from 4 mice, p < 0.05). The high density of muscarinic M_3_ receptors localised to Stratum Lacunosum Moleculare where TA inputs synapse in CA1 (Levey et al., 1995) suggests a role for M_3_ receptors modulating the TA pathway. Supporting a role for M_3_ receptors, the selective M_3_ receptor antagonist DAU5884 (1 µM) (Gosens et al., 2004) prevented the EPSC depression and increase in PPR caused by CCh (Figure 4F-G; EPSC 105 ± 11 %, n = 6 from 4 mice, p > 0.05; PPR 101 ± 6 %, n = 6 from 4 mice, p > 0.05) suggesting that TA pathway synaptic transmission onto CA1 pyramidal neurons is modulated by presynaptically located muscarinic M_3_ receptors.

**Figure 4.**
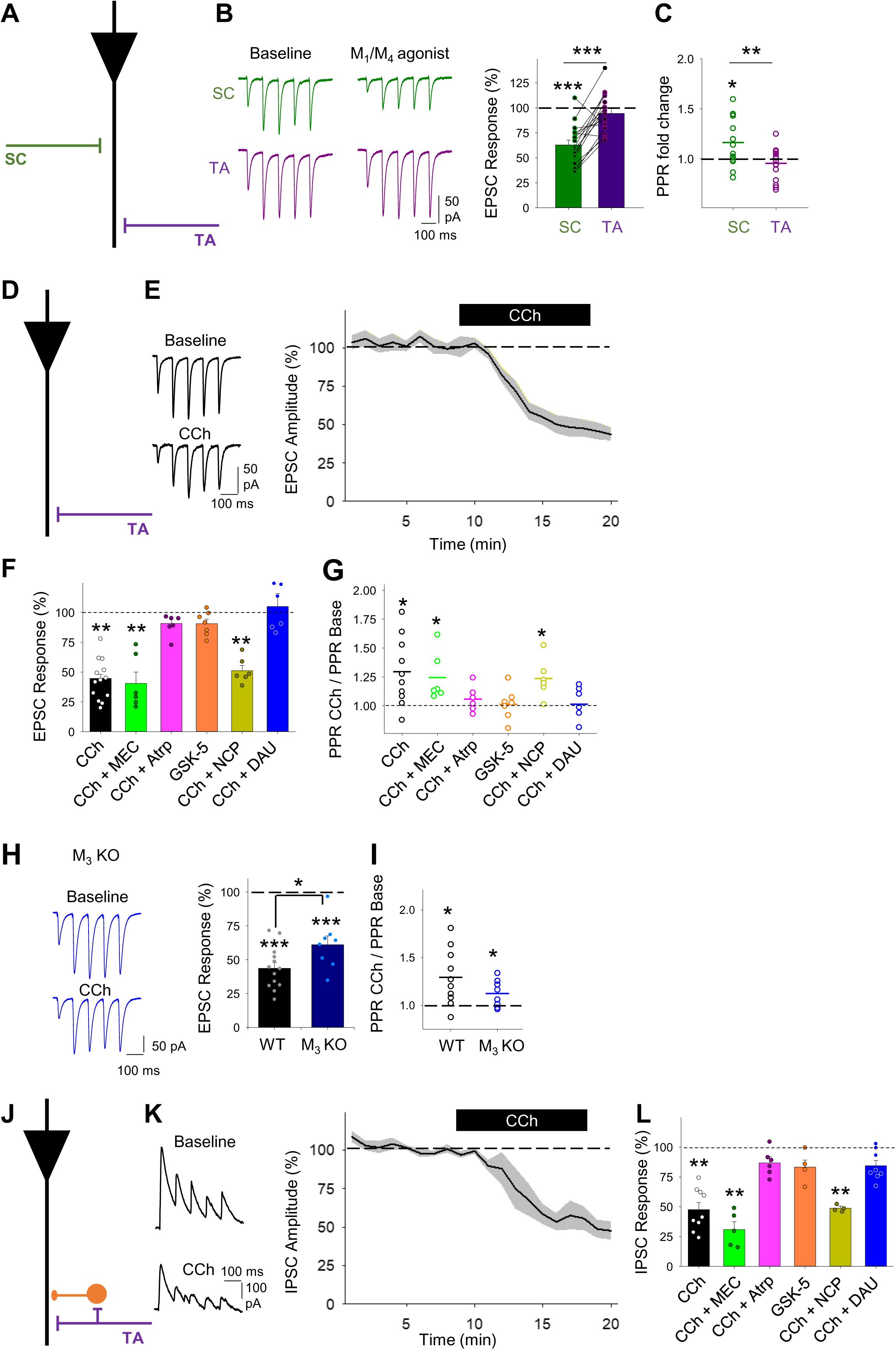
Muscarinic M_3_ receptors modulate temporoammonic pathway EPSCs and disynaptic IPSCs in CA1 pyramidal neurons. **A**, Schematic illustrating recording of pharmacologically isolated EPSCs from temporoammonic (TA) and Schaffer collateral (SC) pathways. **B-C,** The dual M_1_ and M_4_ and muscarinic receptor agonist Compounnd1 (1 µM) depresses evoked EPSCs (B) and increases paired pulse ratio (PPR) (C) for SC (green) but not TA (purple) pathway. **D**, Schematic illustrating recording of pharmacologically isolated EPSCs from temporoammonic (TA) pathway. **E**, CCh (10 µM) reliably reduced evoked EPSC amplitudes. **F-G**, Pharmacology of cholinergic depression of EPSCs. CCh-induced depression (F) is prevented by application of muscarinic receptor antagonist atropine (Atrp, 10 µM) or M_3_ receptor antagonist DAU 5884 (DAU, 1 µM) but not M_1_ receptor antagonist Nitrocaramiphen (NCP, 1 µM) or nicotinic receptor antagonist mecamylamine (MEC, 25 µM) and is not replicated by M_1_ receptor agonist GSK-5 (500 nM). PPR changes reflect conditions of cholinergic-induced EPSC depression (G). **H-I**, Comparison of the effects of CCh on TA pathway-evoked EPSCs in wild type (WT) and M_3_ receptor knockout mice (M_3_ KO). EPSC depression (H) and PPR increase (I) by CCh were reduced in slices from M_3_ KOs in comparison to WT. **J**, Schematic illustrating recording of disynaptic feedforward IPSCs from pyramidal neurons at 0 mV in TA pathway. **K**, CCh (10 µM) reliably reduced evoked IPSC amplitudes. **L**, Pharmacology of cholinergic depression of IPSCs. CCh-induced depression is prevented by application of muscarinic receptor antagonist atropine or M_3_ receptor antagonist DAU 5884 but not M_1_ receptor antagonist Nitrocaramiphen or nicotinic receptor antagonist mecamylamine and is not replicated by M_1_ receptor agonist GSK-5. Data are mean ± SEM; Inter group comparison one-way ANOVA with post hoc Bonferroni correction. Two tailed unpaired Student’s T-test *** p < 0.001 ** p < 0.01 * p < 0.05.

To confirm the involvement of presynaptic muscarinic M_3_ receptors, we tested the effects of CCh in M_3_ receptor knock out mice (M_3_ KO) (Yamada et al., 2001). Although TA evoked EPSCs recorded from M_3_ KO slices were reduced by CCh with an associated increase in PPR (Figure 4H-I; EPSC, 61 ± 6 %, n = 8 from 4 mice, p < 0.001; PPR, 112 ± 5 %, p < 0.05), this CCh-induced depression was less than that recorded in WT slices (Figure 4H; WT EPSC vs M_3_ KO EPSC, p < 0.05). This confirms the pharmacological data for presynaptic M_3_ receptor involvement in the TA pathway but also suggests some compensation for M_3_ receptor deletion within M3 KO mice. The most likely subunit to compensate for M_3_ deletion are M_1_ receptors that are expressed in pyramidal cells and are also coupled to Gq signalling pathways. Therefore, to further explore possible compensatory mechanisms, we tested the selective muscarinic M_1_ receptor agonist GSK-5 in the M_3_ KO mice (Figure S5). M_1_ receptor activation depolarises and increases spike rates in pyramidal neurons (Buchanan et al., 2010) thereby increasing spontaneous EPSCs. Application of GSK-5 increased spontaneous EPSC frequency in slices from both WT and M_3_ KO mice (Figure S5A) but caused a selective decrease in TA EPSC and corresponding increase in PPR in the M_3_ KO but not the WT (Figure S5B-C). This indicates that M_1_ receptors partially replace the deleted M_3_ receptors at presynaptic TA terminals in M_3_ KO mice.

Feedforward synaptic inhibitory transmission in the TA pathway was isolated by holding the membrane voltage at 0 mV (see methods; Figure 4J). As previously described (Figures 1&2), CCh depressed IPSCs without an effect on PPR (Figure 4K-L; IPSC, 48 ± 6 %, n = 9 from 4 mice, p < 0.01; PPR, 108 ± 3 %, n = 9 from 4 mice, p > 0.05). The pharmacological data again supported a role for M_3_ receptors. Nicotinic receptor antagonist mecamylamine (25 µM) did not prevent CCh-induced depression (Figure 4L; 31 ± 6 %, n = 5 from 3 mice, p < 0.01) but the muscarinic receptor antagonist atropine (10 µM) did (Figure 4L; 87 ± 5 %, n = 6 from 3 mice, p > 0.05), demonstrating that, as for excitatory synaptic transmission, inhibitory inputs to CA1 pyramidal neurons are depressed by muscarinic receptor activation. Muscarinic M_1_ receptors did not alter TA IPSC as the agonist GSK-5 was unable to modulate inhibitory synaptic transmission (Figure 4L; GSK-5 500 nM; 83 ± 6 %; n = 4 from 2 mice, p > 0.05) and the M_1_ receptor antagonist nitrocaramiphen was unable to block the CCh effect (Figure 4L; nitrocaramiphen 100 nM, 49 ± 2 %, n = 4 from 2 mice, p < 0.01). Similar to excitatory transmission, muscarinic M_3_ receptor antagonist (DAU5884 1 µM) blocked TA pathway IPSC modulation by CCh (Figure 4L; 84 ± 4 %, n = 8 from 4 mice, p > 0.05). These results show that M_3_ muscarinic receptors are located at presynaptic TA terminals where they depress release of glutamate onto CA1 pyramidal neurons and feedforward inhibition within the TA pathway.

### Cholinergic disinhibition enhances CA1 output in response to temporoammonic but not Schaffer collateral input

The modulation of hippocampal synaptic transmission and in particular the differential regulation of excitatory-inhibitory balance of SC and TA synaptic pathways predicts that acetylcholine prioritises CA1 response to inputs from entorhinal cortex via the TA pathway. To test this prediction, we monitored spike generation in CA1 pyramidal neurons in response to SC and TA pathway stimulation using trains of 10 stimuli at 10 Hz given to SC or TA pathways. The stimulus intensities were set so that post synaptic potentials (PSPs) were suprathreshold for action potential initiation on some but not all stimuli (P_spike_; see methods). Application of 10 µM CCh depolarised CA1 pyramidal neurons (average depolarisation 5.3 ± 0.7 mV) so to dissociate the effects of CCh on membrane potential and synaptic inputs current was initially injected to maintain membrane potential at baseline (i ≠ 0) and assessed changes in spike probability. Subsequently, the injected current was removed (i = 0) to examine how cholinergic depolarisation affected spike probability. With membrane potential maintained at baseline levels, CCh dramatically reduced the probability of spikes generated by SC pathway stimulation (Figure 5A_1_-C_1_; P_spike_ baseline 0.59 ± 0.07 vs CCh i ≠ 0 0.14 ± 0.05, n = 12 from 5 mice, p < 0.001) and required more stimuli within a train and therefore a longer delay to generate the first spike (Figure 5A_1_-C_1_; baseline, 298 ± 58 ms vs CCh i ≠ 0, 775 ± 83 ms, p < 0.001). With current injection removed and membrane potential allowed to depolarise, spike probability increased slightly but failed to return to baseline levels (Figure 5A_1_-C_1_; P_spike_ 0.33 ± 0.05, p < 0.05 baseline versus CCh i = 0). In contrast, CCh application had little effect on TA pathway driven spike probability and delay to the first spike when the membrane potential was maintained at baseline levels (Figure 5A_2_-C_2_; P_spike_ baseline, 0.33 ± 0.06 vs CCh i ≠ 0, 0.43 ± 0.08, n = 15 from 10 mice, p > 0.05; delay to spike baseline, 397 ± 56 ms vs CCh i ≠ 0, 483 ± 66 ms, p > 0.05). However, with current injection removed and CA1 neurons allowed to depolarise spike probability increased and the delay to first spike shortened (Figure 5A_2_-C_2_; P_spike_ 0.61 ± 0.06, p < 0.01 vs baseline; delay to spike 280 ms ± 27 ms, p < 0.05 vs baseline).

**Figure 5.**
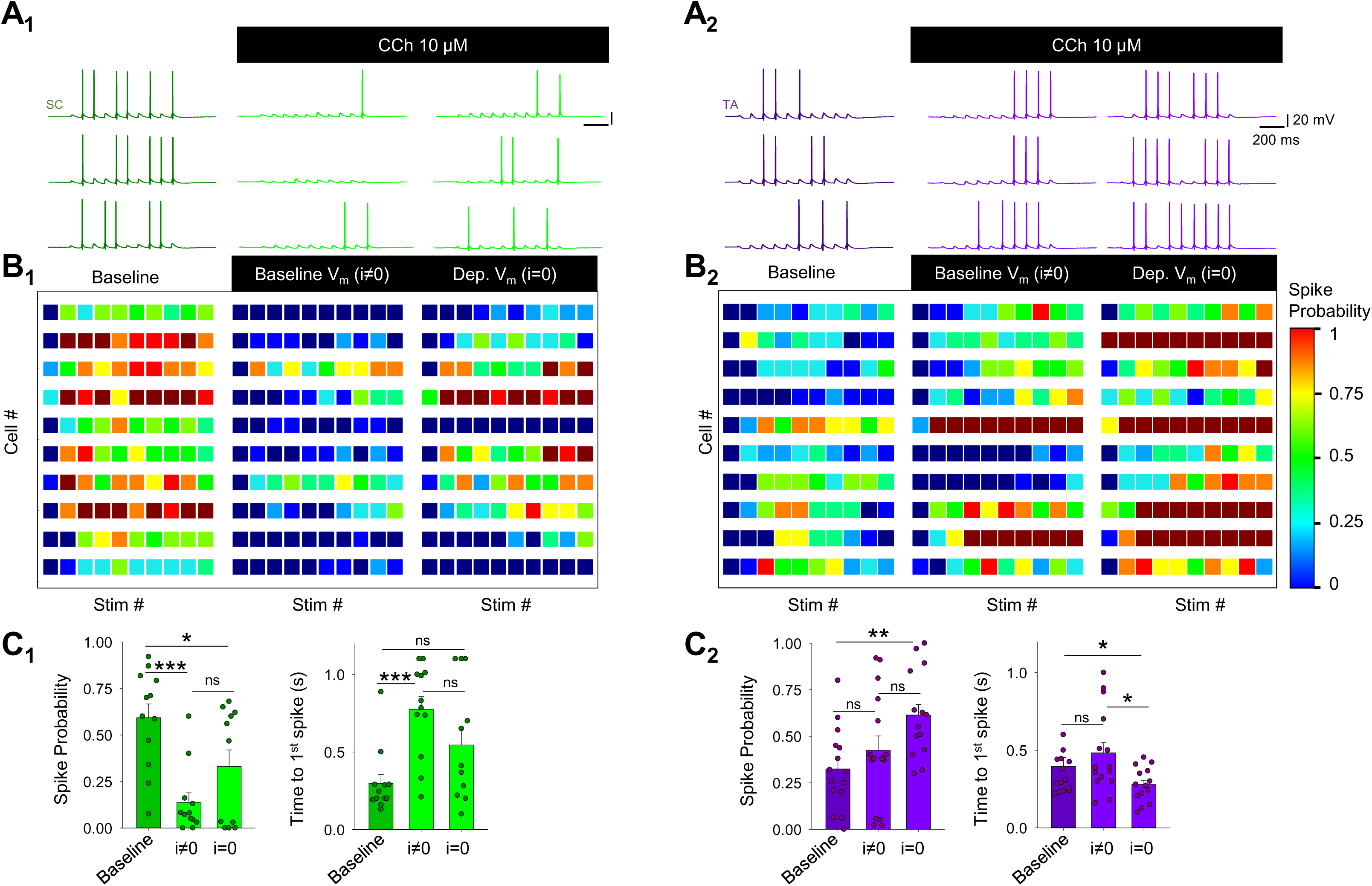
Cholinergic receptor activation enhances CA1 response to temporoammonic over Schaffer collateral input. **A**, Responses in CA1 pyramidal neurons to 10 stimuli at 10 Hz given to Schaffer collateral (SC, A_1_) or temporoammonic (TA, A_2_) input pathways. After application of CCh (10 µM), membrane potential (V_h_) is initially held at baseline levels by injection of current (i≠0) and then allowed to depolarise (i=0). **B**, Heat maps depicting spike probability for 10 stimulation pulses from 10 cells for SC (B_1_) and TA (B_2_) input pathways before and during CCh application. **C**, Spike probability and time to first spike for SC (C_1_) and TA (C_2_) input pathways. Spike probability decreased after CCh application in SC pathway but increased in TA pathway. Data are mean ± SEM; One-way ANOVA with repeated measures and post hoc Bonferroni correction *** p < 0.001 ***p < 0.01 * p < 0.05.

Since CCh or endogenous acetylcholine reduce excitatory synaptic inputs from the SC and TA pathways equally (Figures 1&2), our data suggest the CCh-induced increase in spike probability in response to TA pathway input is caused by a frequency-dependent depression of feedforward inhibition, and therefore increase in excitatory-inhibitory balance, selectively in the TA pathway (Figure 2). Indeed, a substantial hyperpolarising envelope driven by inhibitory synaptic inputs was seen in spike probability recordings from both SC and TA pathways and could be removed by application of a GABA_A_ receptor antagonist (picrotoxin, 50 µM) (Figure 6A-B; SC hyperpolarising envelope −2.45 ± 0.63 mVs, n = 15 from 5 mice versus SC GABA_A_ antagonist −0.10 ± 0.66 mVs, n = 7 from 2 mice, p < 0.05; TA hyperpolarising envelope −3.01 ± 0.46 mVs, n = 23 from 9 mice versus TA GABA_A_ antagonist −1.19 ± 0.52 mVs, n = 7 from 2 mice, p < 0.05). To test the importance of CCh effect on inhibition for prioritisation of TA inputs we next repeated spike probability experiments in the presence of the GABA_A_ receptor antagonist. Under these experimental conditions SC pathway behaved similarly, decreasing spike generation probability upon CCh exposure when membrane potential was kept unaltered (Figure 6C_1_-D_1_; baseline, 0.7 ± 0.06 and CCh i ≠ 0, 0.21 ± 0.06, n = 9 from 3 mice, p < 0.01) and showed an increase during depolarisation without reaching baseline levels (0.44 ± 0.07, p < 0.05 versus baseline), which was correlated with delay to first spike (baseline, 318 ms ± 67 ms, CCh i ≠ 0, 673 ms ± 122 ms, CCh i = 0, 397 ms ± 92 ms, p < 0.05 baseline versus CCh i ≠ 0). In contrast, the TA pathway, which increased P_spike_ after CCh when PSP included both excitatory and inhibitory drive, yielded a similar spike probability outcome to SC pathway when inhibition was blocked, decreasing spike probability whether membrane potential was depolarised or not (Figure 6C_2_-D_2_; baseline, 0.58 ± 0.06; CCh i ≠ 0, 0.17 ± 0.05; CCh i = 0, 0.37 ± 0.08; n = 8 from 4 mice; p < 0.01 baseline versus CCh i ≠ 0 and p < 0.05 baseline versus CCh i = 0). This was associated with increases in the delay to first spike (Figure 6C_2_-D_2_; baseline 246 ms ± 23 ms; CCh i ≠ 0, 631 ms ± 119 ms; CCh i = 0, 464 ms ± 119 ms; p < 0.05 baseline versus CCh i ≠ 0).

**Figure 6.**
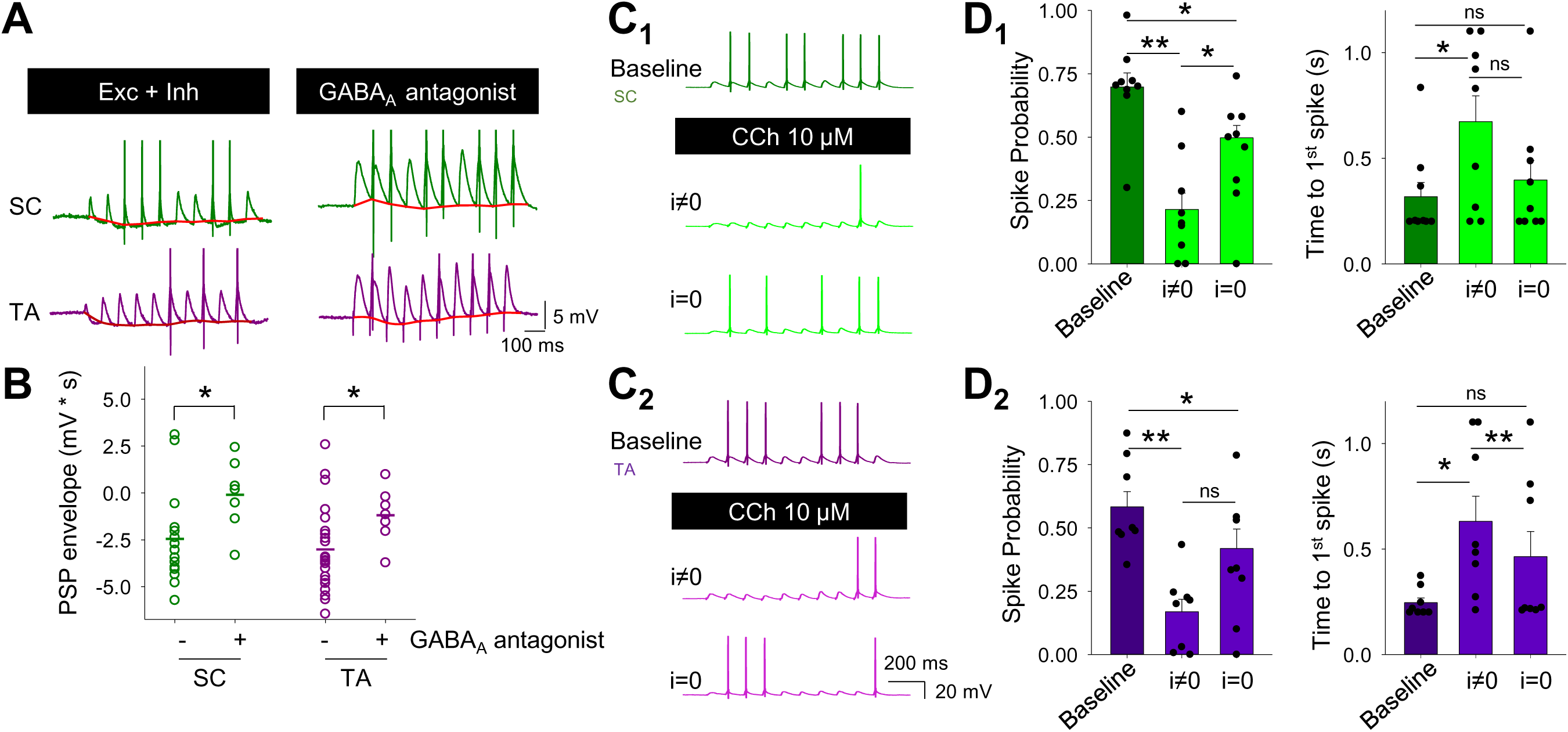
Cholinergic enhancement of CA1 responses to temporoammonic inputs is mediated by inhibition. **A-B**, Inhibition of GABA_A_ receptors with picrotoxin (50 µM) reduced the underlying hyperpolarising envelope (red) in response to 10 stimuli at 10 Hz (D) and increased spike probability for both SC (green) and TA (purple) input pathways to CA1 pyramidal neurons (E). **C-D**, In the presence of GABA_A_ receptor antagonist, CCh reduced spike probability and increased time to spike in both SC (C_1_-D_1_) and TA (C_2_-D_2_) input pathways. Data are mean ± SEM; One-way ANOVA with repeated measures and post hoc Bonferroni correction ** p < 0.01 * p < 0.05.

Finally, we sought to confirm that endogenous release of acetylcholine in the hippocampus also decreases the probability of SC evoked spikes and increases the probability of TA evoked spikes. To test this, we reverted to optogenetic stimulation of cholinergic fibers in mice expressing ChR2 in cholinergic neurons. This produced no change in membrane potential and therefore required no current injection to maintain a constant resting potential. After 5 minutes of 2 Hz light stimulation of acetylcholine release the probability of spiking in response to SC pathway stimulation decreased (Figure 7A-C; SC P_spike_ baseline 0.61 ± 0.04 vs ACh release 0.44 ± 0.06, normalized SC decrease 0.77 ± 0.1, n = 13 from 8 mice, p < 0.05), while probability of spiking in response to TA pathway stimulation increased (Figure 7A-C; TA P_spike_ baseline 0.43 ± 0.05 vs ACh release 0.62 ± 0.05, normalized TA increase 1.71 ± 0.29, n = 14 from 7 mice, p < 0.005). This opposite modulation of SC and TA pathways was striking in a subset of recordings made from both pathways in the same neuron (P_spike_ SC vs TA, n = 11, p < 0.05). These changes were matched by an increase to the time to first spike in the SC pathway and a decrease for the TA pathway (Figure 7D; SC normalised time to first spike 1.77 ± 0.28, p < 0.05 and TA 0.72 ± 0.05, p < 0.001). The effects of light evoked stimulation on CA1 spike probability and delay were completely blocked by the inclusion of muscarinic and nicotinic antagonists (Figure 7A,C,D; SC normalised P_spike_ 1.04 ± 0.1, n = 9 from 4 mice, p > 0.05 & TA 0.82 ± 0.16, n = 6 from 4 mice, p > 0.05; SC normalised time to first spike 0.86 ± 0.06, p > 0.05 & TA 1.33 ± 0.17, p > 0.05). Therefore, endogenous acetylcholine release down-regulates CA1 pyramidal neuron responses to SC pathway and up-regulates responses to TA pathway. Altogether, our data indicate that cholinergic receptor activation produces a decrease of spike output in response to SC activity while enhancing output in response to TA activity via a differential effect on feedforward inhibition to CA1.

**Figure 7.**
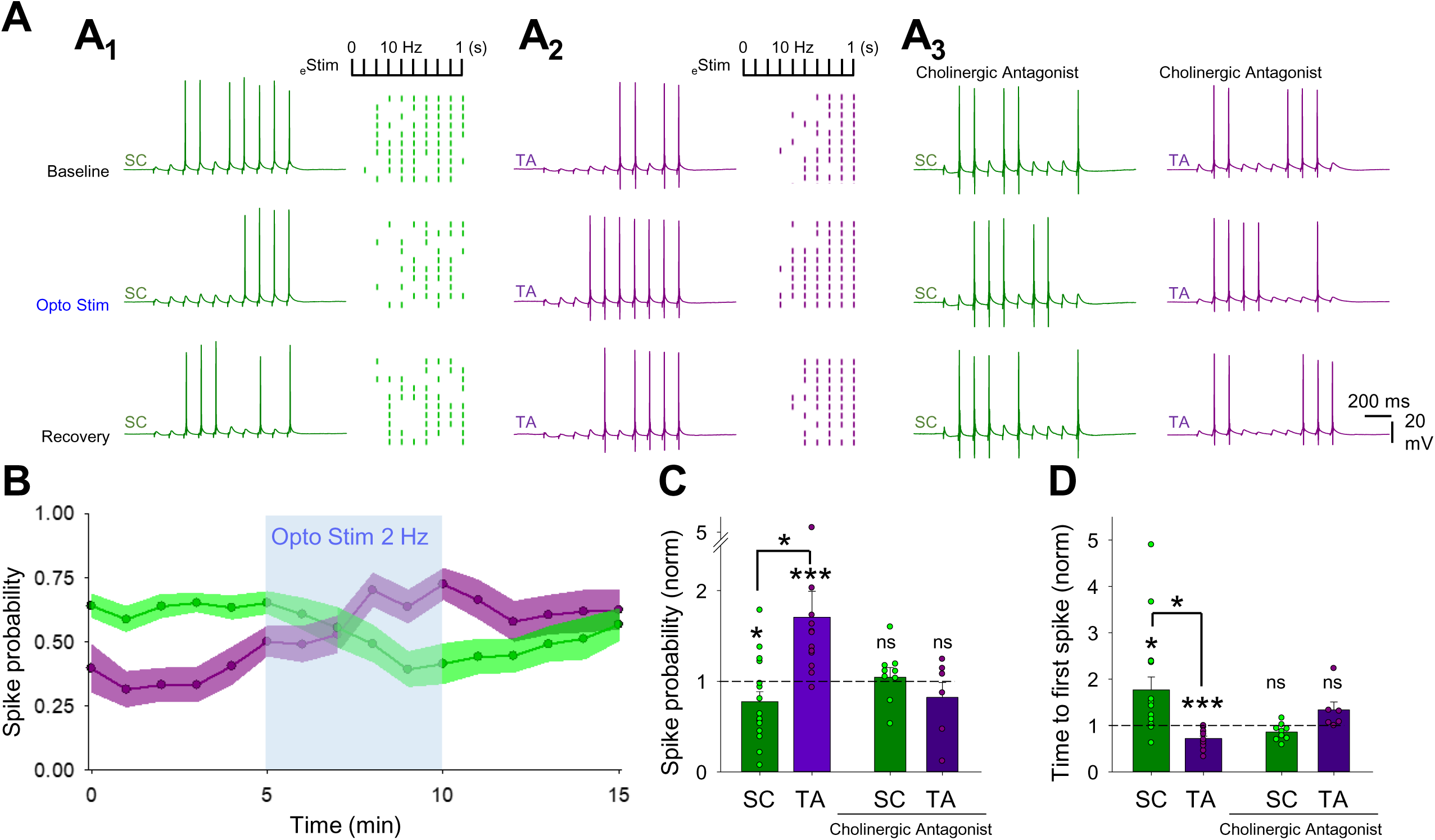
Endogenous acetylcholine release enhances CA1 response to temporoammonic over Schaffer collateral input. **A**, Modulation of SC (A_1_) and TA (A_2_) pathway spike generation by endogenous optogenetically-evoked release of acetylcholine (2 Hz for 5 mins). Raster plots show representative experiments where SC and TA pathway are stimulated alternately with trains of 10 stimuli. Cholinergic receptor antagonists, atropine (25 µM) and mecamylamide (50 µM) prevented modulation of spike generation in SC and TA pathways by endogenous acetylcholine release (A_3-4_). **B**, Time course of spike probability modulation on SC and TA pathway by endogenous release of acetylcholine. **C-D**, Quantification of spike probability (C) and time to first spike (D) on SC and TA pathway either on naïve or in the presence of cholinergic antagonists. Data are mean ± SEM; Inter group comparison one-way ANOVA with post hoc Tukey’s correction. Two tailed paired Student’s T-test *** p < 0.001 * p < 0.05.

## Discussion

A long-standing and influential theory proposes that acetylcholine release in the hippocampus prioritises novel sensory information input to enable incorporation into memory ensembles (Hasselmo, 2006). This theory is based on computational modelling and the observation that SC synaptic inputs are more sensitive than TA inputs to depression caused by exogenous cholinergic agonists (Dannenberg et al., 2017; Hasselmo, 2006; Hasselmo and Schnell, 1994; Hasselmo et al., 1995). In contrast, we show that excitatory synaptic transmission at SC and TA inputs to CA1 are equally depressed by endogenous acetylcholine released in response to optogenetic stimulation (Figure 1). Furthermore, in the absence of inhibition, we show that this results in a dramatic reduction of spike output from CA1 in response to either SC or TA input (Figure 6). However, when we considered the effects of acetylcholine on local inhibitory networks as well as excitatory inputs, we find that acetylcholine depresses feedforward inhibition in the TA pathway more than the SC pathway over the course of a burst of stimuli (Figures 2&3). This results in an overall enhancement of spike output from CA1 in response to TA input but not the SC input (Figures 5&7) supporting the hypothesis that acetylcholine enhances responses to novel sensory information arriving via the TA pathway.

The regulation of local inhibitory networks by acetylcholine is therefore central to prioritisation of TA inputs by acetylcholine and differences in the regulation of synaptic output from interneuron subtypes are a critical factor. Although the interneuron subtypes engaged by the SC and TA pathways are a mixed population, the difference in IPSC kinetics and sensitivity to the µ-opioid receptor agonist DAMGO in our recordings support previous findings that PV^+^ cells form the majority of feedforward inhibition in the SC pathway whereas CCK^+^ cells are the major contributors to feedforward inhibition in the TA pathway (Basu et al., 2013; Freund and Katona, 2007; Glickfeld and Scanziani, 2006; Klausberger and Somogyi, 2008; Milstein et al., 2015). Crucially, the synaptic output from PV^+^ and CCK^+^ interneurons is differentially regulated by acetylcholine. Whilst both outputs are depressed by acetylcholine, the depression of CCK^+^ output is greater over the course of a burst of stimuli showing enhanced depression for later responses in the burst, mirroring the effect of acetylcholine on feedforward inhibition in the TA pathway. Acetylcholine does not cause a greater depression for later responses in the burst for PV^+^ synaptic output and therefore feedforward inhibition in the SC pathway is relatively greater over a burst of stimuli reducing the impact of SC stimulation when acetylcholine is present. Interestingly, the excitability of different interneuron subtypes is regulated by different cholinergic receptors with M_3_ receptors in CCK^+^ interneurons, M_1_ receptors in PV^+^ and NPY^+^ interneurons and nicotinic α2 receptors in OLM feedback interneurons (Cea-del Rio et al., 2011; Cea-del Rio et al., 2010; Leao et al., 2012; Raza et al., 2017; Yi et al., 2014). This further supports the major contribution of CCK^+^ interneurons in the TA pathway since the M_3_ receptor antagonist DAU5884 completely blocked the CCh-induced depression of feedforward inhibition in the TA pathway.

The differential regulation of SC and TA pathways is mediated by selective expression of M_4_ and M_3_ receptors. The targeting of M_3_ and M_4_ receptors to presynaptic terminals of TA and SC axons respectively fits with a broader picture of highly specific localisation of muscarinic receptor subtypes to cellular and subcellular domains within the hippocampus that includes the localisation of M_2_ receptors to inhibitory presynaptic terminals of PV^+^ basket cells. This agrees with the observed highly laminar localisation of M_3_ receptors in the Stratum Lacunosum Moleculare, M_4_ receptors in Stratum Radiatum and M_2_ receptors in the Stratum Pyramidale ((Levey et al., 1995) but see (Goswamee and McQuiston, 2019)). At each terminal, muscarinic receptors depress neurotransmitter release probability (Dasari and Gulledge, 2011; Levey et al., 1995; Szabo et al., 2010; Thorn et al., 2017) and we show that this includes M_3_ receptors targeted to presynaptic terminals of TA axons where they depress release of glutamate. M_3_ receptors are also expressed in CCK^+^ interneurons where they increase excitability (Cea-del Rio et al., 2011; Cea-del Rio et al., 2010) and our data suggest that M_3_ receptors expressed in these cells can also regulate release of GABA at synapses onto pyramidal cells (Figures 3&4). Given the importance of the TA input for synaptic plasticity in the hippocampus (Bittner et al., 2015; Takahashi and Magee, 2009) it is expected that M_3_ receptors play an important role in hippocampal-dependent learning. However, the evidence from studies using mice with genetic deletion of M_3_ receptors is somewhat equivocal (Poulin et al., 2010; Yamada et al., 2001). A potential explanation lies in the compensation for deletion of M_3_ with expression of M_1_ receptors (Figure S4) that couple to similar Gq-mediated signalling pathways and it is interesting that knockin mutations of phosphorylation-deficient M_3_ receptors with potentially less compensation show greater effects on learning and memory (Poulin et al., 2010). The compensation for M_3_ deletion by M_1_ receptors is somewhat surprising since M_1_ receptors are generally expressed widely in somatic and dendritic cellular domains in pyramidal cells and interneurons where they regulate intrinsic excitability leading to effects on synaptic plasticity and network oscillations (Atherton et al., 2016; Betterton et al., 2017; Buchanan et al., 2010; Dennis et al., 2016; Fisahn et al., 2002; Levey et al., 1995; Mitsushima et al., 2013; Shinoe et al., 2005; Tigaret et al., 2018) but are not generally found in presynaptic terminals (Yamasaki et al., 2010).

Cholinergic neurons in vivo fire at frequencies ranging from 0.3 – 5 Hz with higher frequencies recorded during waking activity (Hangya et al., 2015; Simon et al., 2006). Responses to salient events such as positive or negative reinforcement have been demonstrated (Hangya et al., 2015; Lovett-Barron et al., 2014; Teles-Grilo Ruivo et al., 2017), but even in these conditions cholinergic firing rates do not increase dramatically but rather activity across cholinergic neurons is synchronised (Hangya et al., 2015). Interestingly, release of acetylcholine plateaus at firing rates around 2 Hz (Jing et al., 2018) indicating that the dynamic range of acetylcholine release occurs at frequencies below 2 Hz. Therefore, optogenetic stimulation that synchronises release at 2 Hz over extended time periods is likely to be physiologically maximal. Cholinergic neurons are also reported to co-release glutamate and more prominently GABA both from long-range projections and also local cholinergic interneurons (Saunders et al., 2015; Takacs et al., 2018; Yi et al., 2015). However, we found no evidence for glutamate or GABA release after optogenetic stimulation of cholinergic fibres (Figure 1D). Therefore, under our experimental conditions, optogenetic stimulation of cholinergic fibres at 2 Hz likely provides a maximally effective release of acetylcholine without co-release of glutamate or GABA that was mimicked by exogenous application of 10 µM CCh.

Acetylcholine increases the output gain from primary sensory cortices enhancing signal-to-noise for new sensory information and desynchronising the local cortical network by reorganising inhibition to disinhibit pyramidal neurons (Eggermann et al., 2014; Fu et al., 2014; Letzkus et al., 2011). A contrary situation is reported in the hippocampus where cholinergic activation of dendritically targeting interneurons inhibits pyramidal neurons and potentially gates CA1 output (Haam et al., 2018; Leao et al., 2012; Lovett-Barron et al., 2014). Both of these mechanisms may be important for learning new representations, however, neither of these situations addresses whether acetylcholine prioritises one set of inputs over another. Here, we reveal a novel mechanism whereby acetylcholine alters the short-term dynamics of information processing in CA1 by acting on two distinct muscarinic receptor subtypes located in the SC and TA pathway. It will be interesting to discern in future how these various mechanisms interact across different behavioural epochs.

Multiple compounds have been developed to selectively target M_1_ and M_4_ muscarinic receptors for potential cognitive enhancement whereas M_2_ and M_3_ receptors have received much less attention due to complications with peripheral effects on cardiac and enteric function. The M_1_/M_4_ receptor dual agonist Xanomeline has cognitive enhancing and antipsychotic efficacy in clinical trials (Bodick et al., 1997; Shekhar et al., 2008) and whilst it is not clear whether M_1_ or M_4_ receptors are the key target, in separate studies selective M_1_ agonists and M_4_ agonists have been shown to have memory enhancing and/or antipsychotic efficacy (Chan et al., 2008; Nathan et al., 2013) whereas deletion of M_1_ receptors in mice causes memory deficits (Anagnostaras et al., 2003). Our data provide a mechanism for the actions of M_1_/M_4_ receptor dual agonists such as Xanomeline and Compound1 where activation of M_1_ receptors facilitates synaptic plasticity (Buchanan et al., 2010) and activation of M_4_ receptors prioritises new information to incorporate into memory. Our data also predict that selective activation of M_3_ receptors could potentially facilitate the consolidation of memory by reducing interference from new information. Interestingly, the link that we demonstrate between selective muscarinic receptor activation and distinct interneuron subtypes suggests a mechanism to selectively target and regulate these interneuron populations. This could have therapeutic value in disorders with disruption to specific interneuron populations such as PV^+^ neurons in schizophrenia (Lewis et al., 2005). Overall, acetylcholine release in the hippocampus supports cognition and the identification of specific roles for each muscarinic receptor subtype provides mechanisms to selectively modulate individual aspects of acetylcholine’s actions. The identification of M_3_ receptors as regulators of TA inputs in contrast to M_4_ receptors acting on SC inputs provides a novel mechanism by which specific targeting of these muscarinic receptors could represent a therapeutic strategy to bias hippocampal processing and enhance cognitive flexibility.

## Methods

### Animal Strains

All experiments were performed using male mice. C57BL/6J (Charles River) mice were used as the background strain. The generation of the M_3_ receptor KO mice has been described (Yamada et al., 2001). The M_3_ KO mice used for this study had been backcrossed for 10 times onto the C57BL/6NTac background. Cre reporter allele mice (The Jackson Laboratory) were used to tag specific neuronal populations: Cholinergic neurons (Chat-IRES-Cre; Stock No. 006410), parvalbumin interneurons (B6 PV^CRE^; Stock No. 017320) and cholecystokinin interneurons (CCK-IRES-Cre; Stock No. 012706). Homozygous cre reporter mice were crossed with homozygous Ai32 mice (B6.Cg-Gt(ROSA)26Sortm32(CAG-COP4*H134R/EYFP)Hze/J; Stock No. 024109) to generate litters of heterozygous offspring expressing ChR2.

### Slice preparations

All animal procedures were performed in accordance with Home Office guidelines as stated in the United Kingdom Animals (Scientific Procedures) Act 1986 and EU Directive 2010/63/EU 2010 and experimental protocols were approved by the British National Committee for Ethics in Animal Research.

Brain slices were prepared from P30-40 male mice. Following cervical dislocation and decapitation, brains were removed and sliced in ice-cold sucrose solution containing (in mM): 252 sucrose, 10 glucose, 26.2 NaHCO3, 1.25 NaH2PO4, 2.5 KCl, 5 MgCl2 and 1 CaCl2 saturated with 95% O2 and 5% CO2. Parasagittal slices 350 µm thick were cut using a VT1200 (Leica) vibratome. Slices were transferred to warm (32 °C) aCSF for 30 minutes containing (in mM): 119 NaCl, 10 glucose, 26.2 NaHCO3, 1.25 NaH2PO4, 2.5 KCl, 1.3 MgSO4 and 2.5 CaCl2 saturated with 95% O2 and 5% CO2 and then kept at room temperature until use.

### Electrophysiology

Whole-cell patch clamp recordings were made from hippocampal CA1 pyramidal neurons visualised under infrared differential interface contrast on SliceScope Pro 6000 system (Scientifica). Slices were continually perfused with aCSF at 4-5 ml/min. Patch electrodes (4-7 MΩ resistance) were pulled from borosilicate glass capillaries (Harvard Apparatus) using a PC-87 Micropipette puller (Sutter Instrument). Recording pipettes were filled with either voltage-clamp internal solution (in mM: 117 CsMeSO3, 9 NaCl, 10 HEPES, 10 TEA, 2 MgATP, 0.3 NaGTP, 1 QX-314, 0.3 EGTA at pH 7.3 and 290 mOsm) or current-clamp internal solution (in mM: 135 K-Gluconate, 10 HEPES, 7 glucose, 8 NaCl, 2MgATP, 0.3 NaGTP, 0.2 EGTA at pH 7.3 and 290 mOsm). Electrophysiological recordings were made with an Axoclamp 200B (Molecular Devices) filtered at 5 kHz and digitized at 10 kHz using a CED micro 1401 MKII board and Signal5 acquisition software (Cambridge Electronic Design). Series and input resistances were monitored by applying a 20 pA and 500 ms square pulse. Experiments were neurons displayed >25% change in series resistance were discarded from subsequent analysis. Membrane potentials were not corrected for junction potentials.

*Dual pathway (SC and TA) stimulation*. Bipolar stimulating electrodes were placed in CA3 to stimulate SC fibres and in the Stratum Lacunosum Moleculare (SLM) of subiculum to stimulate TA fibres. Synaptic responses were evoked alternately in either pathway at15 sec intervals. Monosynaptic EPSCs were recorded either at −65 mV membrane potential in the presence of GABA_A_ receptor blocker picrotoxin (50 µM) or in control aCSF at the experimentally determined reversal potential for GABA_A_ receptors (−60 mV). Disynaptic IPSCs were recorded in control aCSF at experimentally determined reversal potential for AMPA receptors (0 mV). NBQX (20 µM) was applied at the end of experiments to ascertain the contribution of direct stimulation of local interneurons to IPSCs and only responses which showed > 70% reduction in IPSCs were used for analysis. In the experiments specified, SC and TA pathway EPSCs and IPSCs were recorded from the same CA1 pyramidal neuron to calculate excitation-inhibition ratio (E-I ratio). EPSC and IPSC contributions were measured as charge transferred by calculating the area of each synaptic response in pC and the ratio of EPSC and IPSC charge for each response determined the E-I ratio. PPRs were calculated by normalising the amplitude of each response to the first response. TA over SC E-I ratio was calculated for each cell before averaging across cells.

*Current clamp experiments* were performed at resting membrane voltage (−61.3 ± 3.5 mV). TA and SC pathways were stimulated at intervals of 20 s with trains of 10 stimuli at 10 Hz. Stimulation intensities were set to generate target spike probabilities between 30-70 %. Spike probability was calculated as the number of spikes/number of stimuli. Time to first spike was measured from the first stimulus in the train. Post synaptic potential (PSP) envelope was measured by calculating the area under the curve generated by joining the points of maximum hyperpolarisation in response to each stimulation as described previously (Chamberlain et al., 2013). Carbachol (CCh 10 µM) -induced depolarisations were neutralised by current injections to maintain a constant membrane voltage (i≠0). To investigate the impact of CCh induced depolarisation, the injected current was removed (i=0).

### Optogenetic stimulation

Blue light from a 470 nm LED was targeted to slices via a 469 nm emission filter, a GFP dichroic mirror (Thorlabs) and the 4x (ChAT-Ai32) or 40x (PV-Ai32 or CCK-Ai32) microscope objective. 5 ms light pulses at 7-9 mW/mm^2^ intensity were used for all stimuli. Optogenetically-evoked IPSCs were recorded from pyramidal neurons at 0 mV membrane potential in the presence of the AMPA and NMDA receptor antagonists NBQX (10µM) and DAPV (50µM).

### Confocal imaging

Recorded slices were permeabilized with 0.1% Triton X-100 (Sigma) and incubated with Alexa avidin (488 nm or 594 nm; ThermoFisher). CA1 pyramidal neurons from Chat-Ai32 mice were labelled with Alexa-594 and test proximity to cholinergic axons using Chat-Ai32 YFP fluorescence.

### Statistical analysis

Experimental unit was defined as cell for all conditions and only one cell recorded from each slice. Cell and animal numbers are reported for all experiments. All data were plotted as the mean ± SEM. Where comparisons between two conditions were made paired or unpaired two-tailed Student’s t-tests were applied as appropriate. For comparisons between more than 2 conditions one-way repeated measures ANOVA tests with Bonferroni post hoc correction were used. The level of significance was set to 0.05 and p values are shown as follows: * P < 0.05; ** P < 0.01; *** P < 0.001. Experiments on WT and M_3_ KO mice were performed blind to genotype.

### Reagents

Carbachol (CCh), NBQX, DCG-IV, D-APV, picrotoxin, atropine, mecamylamine, nitrocaramiphen, DAU-5884 were purchased from Tocris (UK). GSK-5 was synthetized in-house at Eli Lilly and Co. Stock solutions of these compounds were made by dissolving in water. The selective muscarinic M_1_ & M_4_ receptor agonist Compound1 was synthetized in house at Sosei Heptares and dissolved in DMSO for stock solution. The purity of the final compounds was determined by HPLC or LC/MS analysis to be >95%. Additional experimental details relating to the synthesis of Compound1 and associated structures is described in detail in WO2015/118342 which relates to the invention of agonists of the muscarinic M_1_ receptor and/or M_4_ receptor and which are useful in the treatment of muscarinic M_1_/M_4_ receptor mediated diseases.

## Acknowledgements

We thank Paul Anastasiades and Paul Chadderton for critical input to previous versions of the manuscript and all members of the Mellor group for discussion. We also thank Dr. Jürgen Wess (NIH, NIDDK) for providing the M_3_ receptor KO mice This work was supported by Wellcome Trust, Biotechnology and Biological Sciences Research Council (BBSRC).

## Contributions

Conceptualization, J.P-F. and J.R.M.; Methodology, J.P-F. and M.U.; Investigation, J.P-F. and M.U; Provision of reagents, G.A.B., B.G.T., M.S.C., P.J.N. and A.J.H.B.; Visualization, J.P-F., M.U. and J.R.M.; Writing – Original Draft, J.P-F., M.U., and J.R.M.; Writing –Review & Editing, J.P-F., M.U., P.J.N., A.J.H.B. and J.R.M.; Funding Acquisition, J.R.M.; Supervision, J.R.M.

---

**Figure S1.**
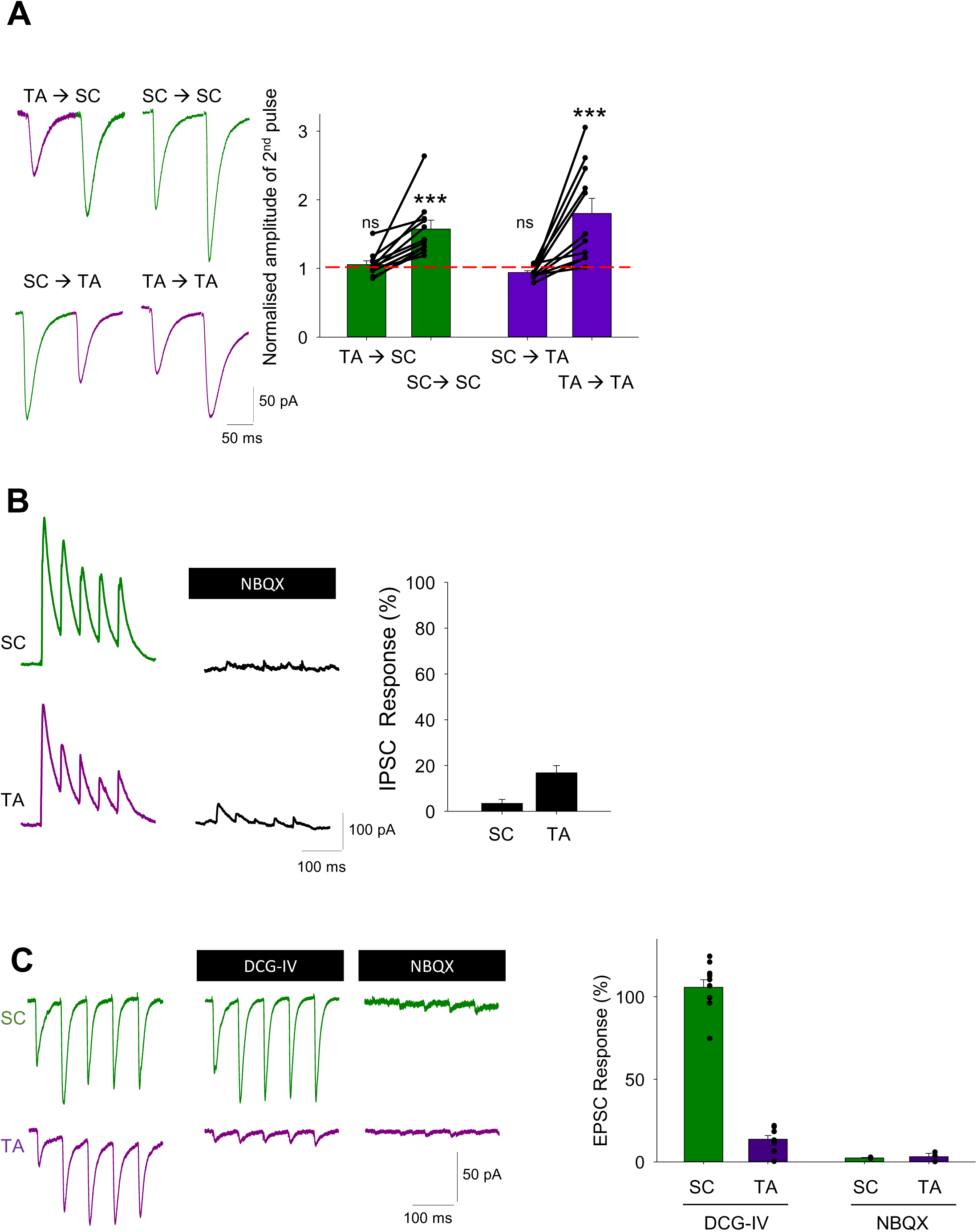
**A**, Independence of SC (green) and TA (purple) pathways was evaluated by the lack of facilitation of the second EPSC response when the alternate pathway was stimulated. **B**, Feedforward IPSCs from SC and TA pathways were recorded at 0 mV and confirmed to be disynaptic by sensitivity to NBQX. **C**, DCG-IV (3 µM) blocked TA pathway but not SC pathway synaptic responses. Application of AMPA receptor antagonist (NBQX 20 µM) blocked responses in both pathways. Data are mean ± SEM; Two tailed unpaired Student’s T-Test. ***P< 0.005.

**Figure S2.**
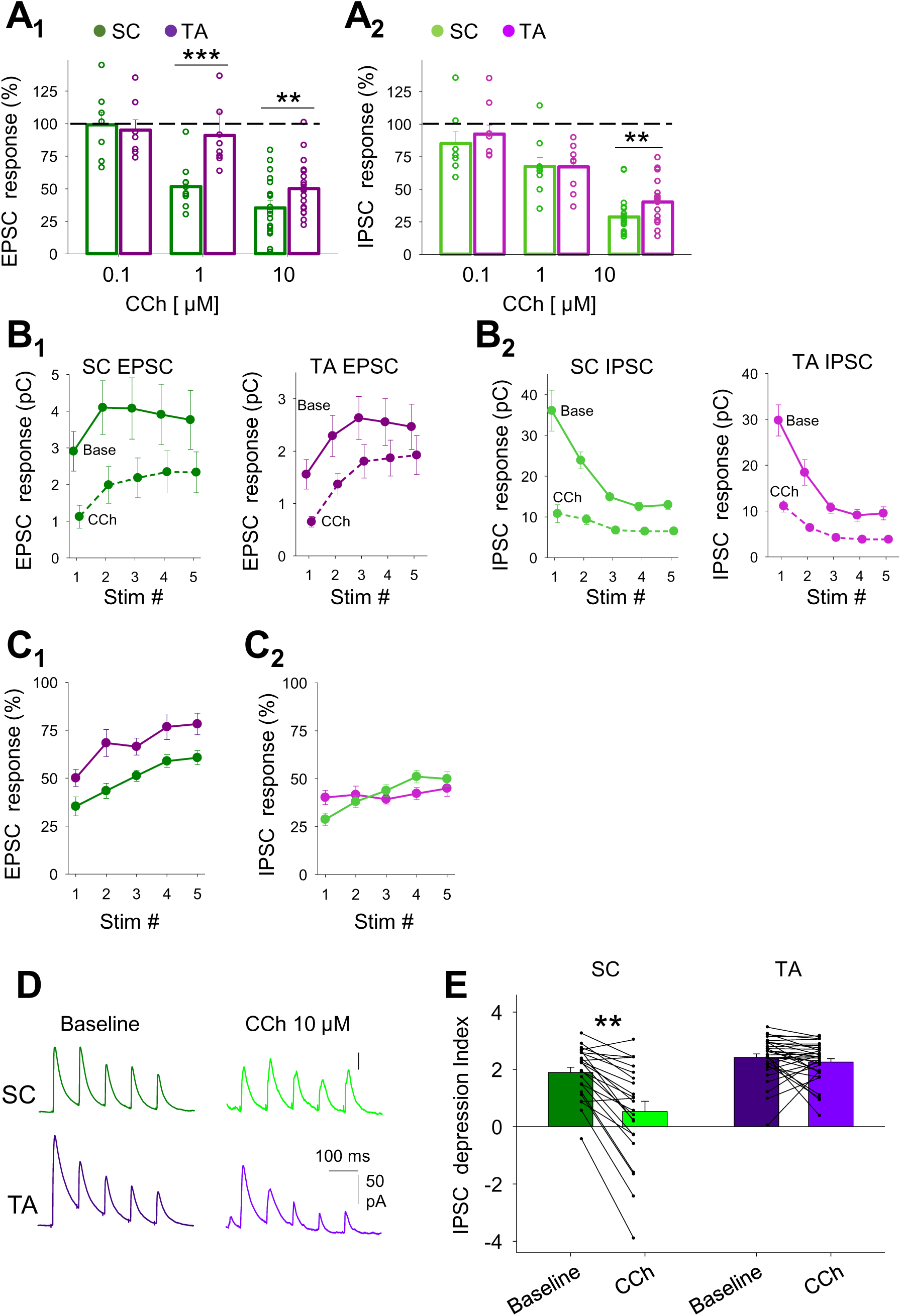
**A**, Dose-response for CCh depression of EPSCs (A_1_) and IPSCs (A_2_) for SC (green) and TA (purple) pathways. **B,** Quantification of SC and TA EPSC (B_1-2_) and IPSC (B_3-4_) charge transfer for each response in the train illustrated in Figure 2A before and after CCh (10 µM) application. **C**, EPSC (C_1_) and IPSC (C_2_) reduction by CCh for each of the 5 stimuli for SC (green) and TA (purple) pathways shown in Figure 2A. **D-E**, CCh reduced the depression index for SC (green) but not TA (purple) disynaptic feedforward IPSCs. Depression index is calculated as the amount of cumulative depression between the 2^nd^ and 5^th^ responses within the train of 5 responses. Data are mean ± SEM; A compared via two tailed unpaired Student’s T-test and E via one-way ANOVA with post hoc Bonferroni correction *** p < 0.001 ** p < 0.01.

**Figure S3.**
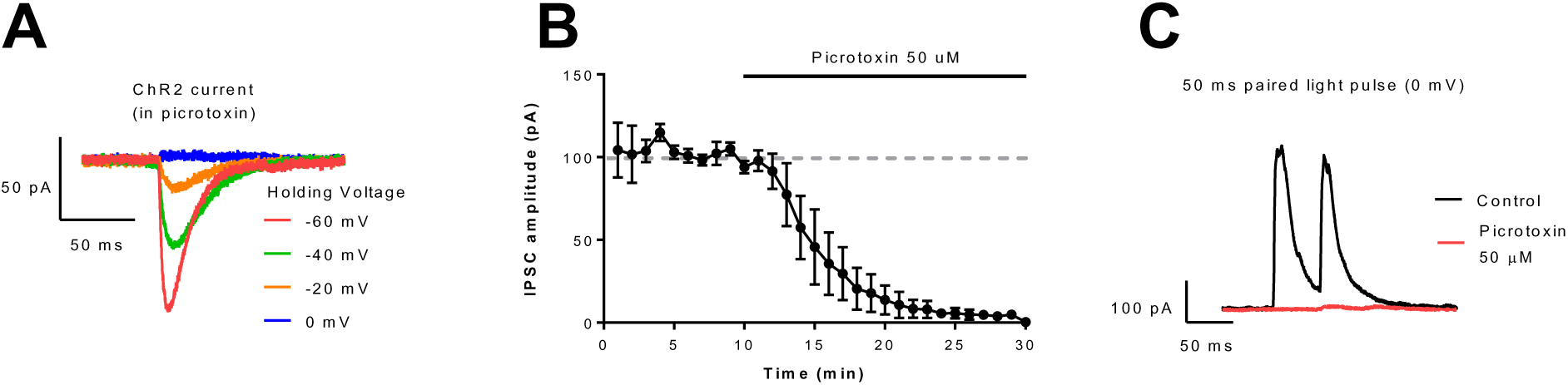
**A**, ChR2 currents at different holding potentials recorded from a CCK^+^ and ChR2 expressing pyramidal neuron in response to 2ms light pulses in the presence of picrotoxin (50 µM). At 0mV (the reversal potential for ChR2) no ChR2 currents are observed. **B-C**, Light evoked GABAergic responses recorded from pyramidal neurons held at 0mV in the presence of NBQX and DAPV are abolished by picrotoxin (50 µM).

**Figure S4.**
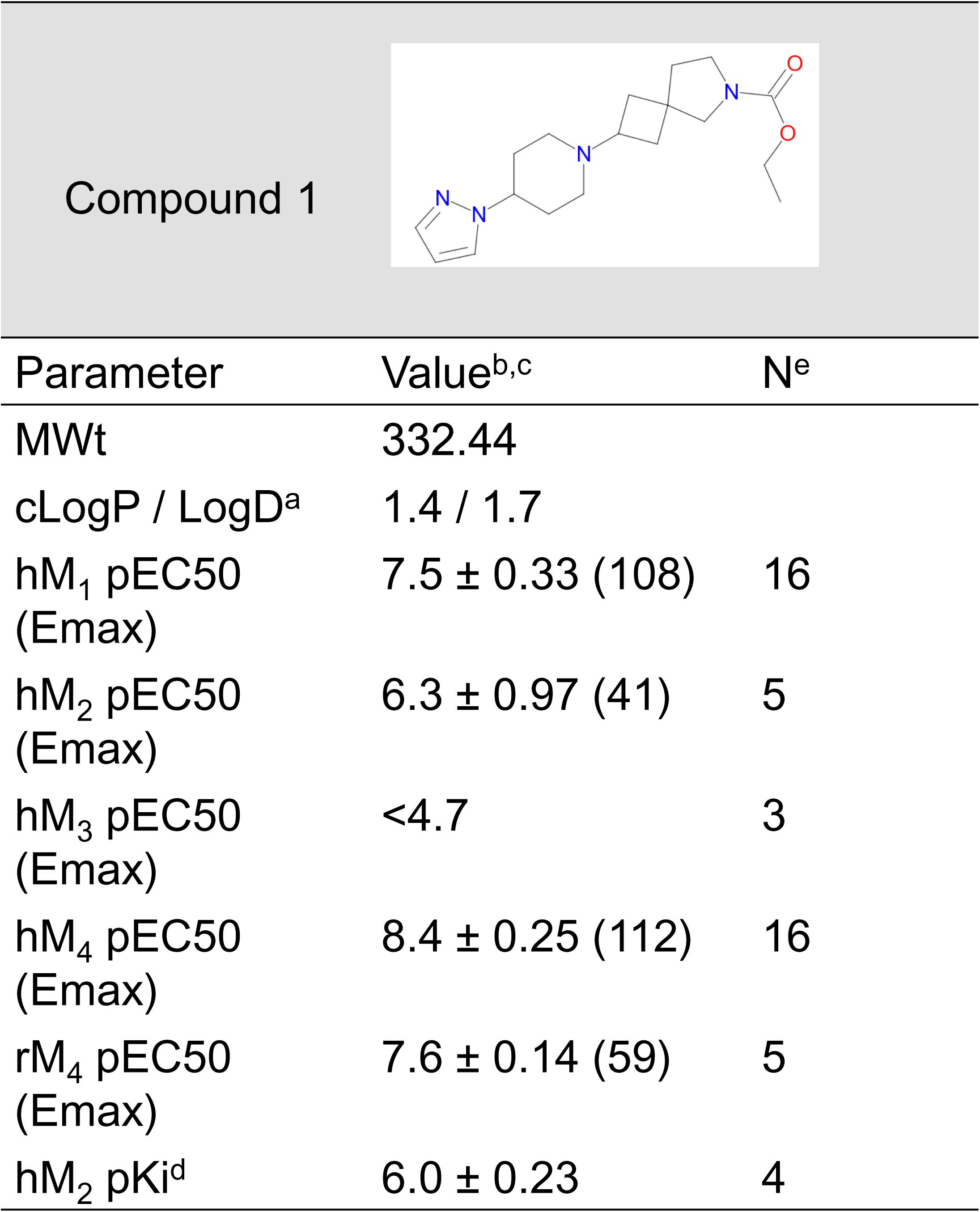
Structure and in vitro pharmacological profile of Compound1. CHO-K1 cells stably expressing the human M_1_–M_4_ and rat M_4_ receptors were used to determine the pharmacological profile of Compound1. ^a^ Calculated LogP value, LogD was measured at pH7.4. ^b^ Compound pEC50 values were measured using phosphor-ERK format (CisBio). Values reported as <4.7 were considered inactive and did not induce a >10% increase in the response at the highest concentration tested (30µM). ^c^ The maximum efficacy (Emax values) are expressed as a percentage of the response of a saturating concentration of acetylcholine (1µM) run in the same assay. ^d^ [3H]-NMS competition binding studies were used to define the affinity (pKi) for Compounnd1 at the human muscarinic M_2_ receptor. ^e^ number of replicates. Data are the mean ± S.E.M. Compound1 can be found within WO2015/118342 which relates to the invention of agonists of the muscarinic M_1_ receptor and/or M_4_ receptor and which are useful in the treatment of muscarinic M_1_/M_4_ receptor mediated diseases.

**Figure S5.**
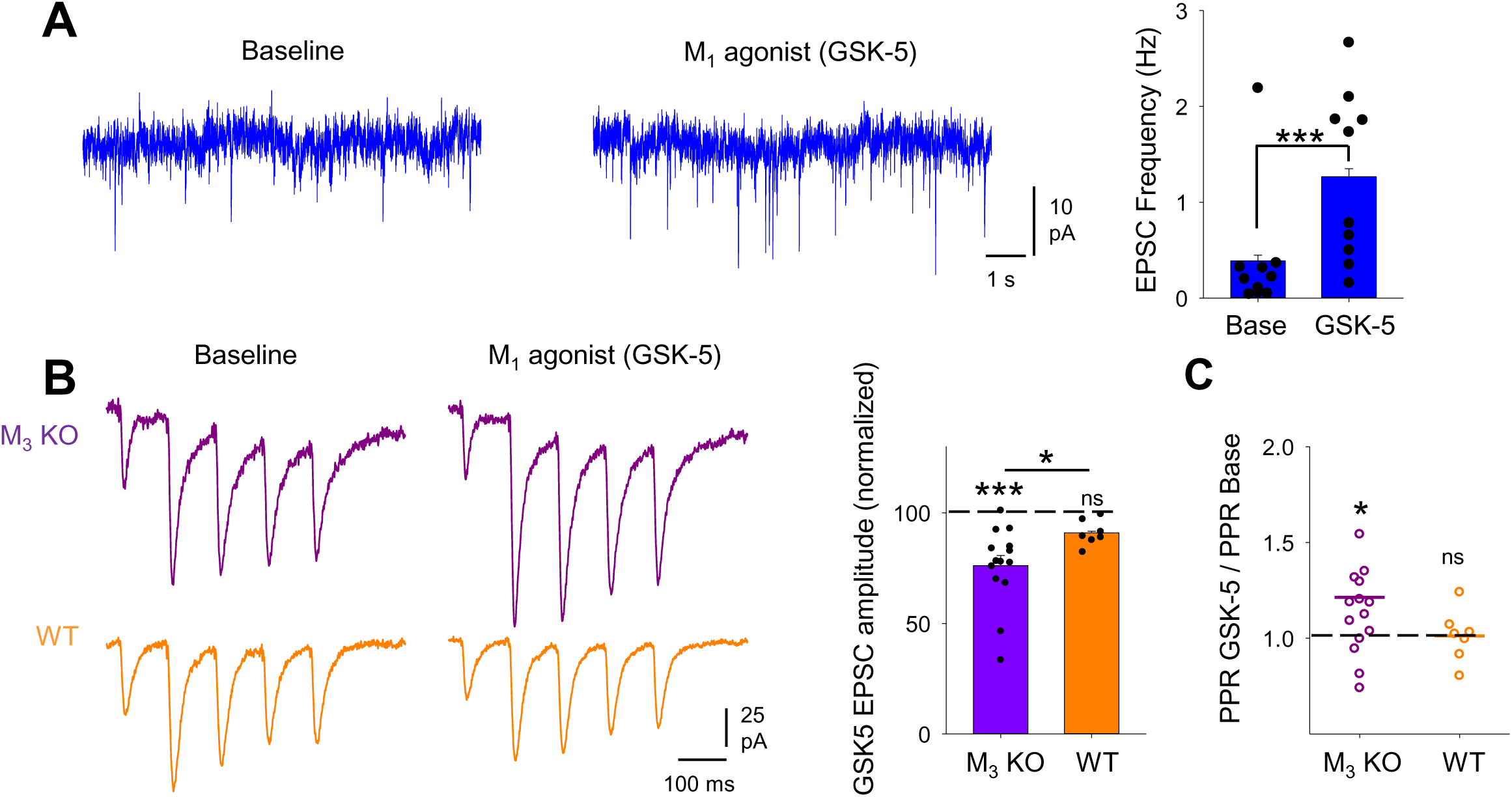
**A**, Muscarinic M_1_ receptor agonist (GSK-5, 500nM) produced an increase in the frequency of spontaneus excitatory events recorded from CA1 pyramidal neurons. **B-C**, GSK-5 caused a reduction of TA pathway EPSC (B) and an increase of PPR (C) in slices from M_3_ KO mice but not in slices from WT mice. Data are mean ± SEM; Inter group comparison one-way ANOVA with post hoc Bonferroni correction. Two tailed paired Student’s T-test *** p < 0.001 * p < 0.05.

